# An Amygdalar Oscillatory Switch Governs Valence Assignment

**DOI:** 10.64898/2025.12.10.693439

**Authors:** E. Teboul, G.L. Weiss, K.A. Amaya, B.T. Stone, P. Antonoudiou, D. Teboul, E.M. Coleman, C. Urzua, J.G. Tasker, J.L. Maguire

## Abstract

Reward pursuit and punishment avoidance are among the most fundamental behaviors necessary for survival. The binary valuation of an experience as either positive or negative - “valence assignment” - is imperative for successful navigation of a regularly updating environment. Mounting evidence highlights the critical role of valence responsive basolateral amygdala (BLA) ensembles in coding valence information. However, how BLA ensembles are recruited to drive real-time valence assignment remains elusive. Here, we show locus coeruleus (LC)-derived norepinephrine coordinates this neural computational process *via* modulatory control over network-organizing BLA parvalbumin-expressing (PV) interneuron activity. Specifically, optogenetic activation of LC to BLA noradrenergic terminals (LC-BLA^NE^) drives real-time negative valence assignment and suppression of BLA fast gamma oscillatory activity *via* BLA interneuronal α1a adrenergic receptor signaling. Conversely, positive valence assignment also requires BLA PV interneuron activity but is associated with an enhancement of local fast gamma power. Together, these converging data highlight a PV-driven amygdalar oscillatory switch that governs valence assignment.

## Introduction

Towards the fundamental goal of all organisms to seek rewards and avoid punishment, animals developed the ability to assign a binary emotional valuation to an experience in real time. In other words, this process, termed “valence assignment,” assigns an animal’s perceived “goodness” or “badness” of a stimulus/stimuli to its experience, irrespective of its scalar value (i.e., emotional salience/magnitude)^1^. Appropriate execution of this process supports learning processes underlying successful navigation of a regularly updating environment. However, inappropriate coordination of this process may result in aberrant responses to innocuous, rewarding, or threatening stimuli, endangering survival. Improper valence assignment has been observed in - and is considered a key component of - several human psychopathologies, including post-traumatic stress disorder^2^, major depressive disorder^3^, suicidality^4,5^, and generalized anxiety disorder^6^; for review, see Antonoudiou et al. (2024)^7^.

Efforts uncovering biological mechanisms supporting this process date back to the late 19th and mid-20th centuries when researchers discovered bilateral temporal lobe ablation, eventually spatially isolating the lesion to the amygdaloid complex, produced impairments in the ability to recognize stimulus positivity or negativity^8–10^. Recent advances highlight the BLA as a central hub in emotion-processing networks, essential for binary emotional valuation across species^11–19^. Collectively, this body of work demonstrates that the BLA, in part, serves to transform diverse inputs into valence-specific outputs that guide appropriate downstream engagement. Notably, spatially intermingled genetically and projection-defined BLA output neuron subtypes coding valence assignment have been described^13,16,17,20,21^, with growing recognition of how both projection targets and molecular expression profiles contribute to establishing valence-specific cellular identities. However, while BLA ensembles and output architecture supporting valence assignment has received considerable attention^12,13,15–18,20–22^, we still lack an understanding of the local computational principles governing the selection and activation of BLA valence-specific ensembles and downstream circuits - a critical step towards developing optimal interventional strategies for psychopathologies associated with aberrant valence assignment.

To this end, neuromodulatory systems with inputs into the BLA have been demonstrated to bidirectionally modulate valence assignment^23–28^, although the cellular mechanisms through which neuromodulators specify valence assignment remain poorly understood. For example, locus coeruleus-derived (LC) noradrenergic transmission in the BLA has been shown to facilitate valence assignment^24–27^. However, while the noradrenergic receptor types and targeted microcircuits involved in the conditioned aspect of valence coding have been described^24–26^, the mechanisms through which noradrenergic signaling in the BLA governs real-time valence assignment have been elusive, though evidence critically implicates alpha 1 adrenergic (α1), but not beta adrenergic receptor activation^26^.

Previous work from our group has demonstrated that BLA α1A receptors, the dominant α1 receptor subtype in the region^29^, are expressed exclusively on interneurons^30^, suggesting that interneurons may play a critical role in the ability of neuromodulators to influence valence assignment. In fact, BLA parvalbumin-expressing (PV) interneurons have been shown to play a critical role in valence-associated behaviors *via* their roles as orchestrators of local population-level activity^30–32^. By shaping the timing and synchrony of local inhibition, noradrenergic input may coordinate BLA output dynamics necessary for appropriate valence coding in real-time. Accordingly, we propose that noradrenergic signaling in PV interneurons facilitates real-time valence assignment by driving specific network states in the BLA. Some evidence exists for this hypothesis, as rhythmic, synchronous neural patterning underlying oscillatory local field potentials (LFPs) can promote activation or maintenance of discrete, content-specific cell ensembles^33–35^ and behavioral states^32,36–41^, potentially *via* facilitation of multiplicative gain and decreased spike variability locally among target neurons^42–44^. In fact, we and others have demonstrated that artificially imposing distinct local oscillatory network states *via* optogenetic control over network-organizing fast-spiking GABAergic/PV interneurons facilitates neural computations underlying real-time perceptual success^45^, spatial memory consolidation^46^, bidirectional fear^37^, reward-seeking^41^, and stress behaviors^32^, among others.

We have previously shown that BLA α1A receptors shape local 40-120 Hz oscillatory network states, principal neuron activity, and affective behavioral state transitioning *via* facilitation of distinct action potential bursting modes among BLA PV interneurons - effects solely attributed to α1A, and not α1B or α1D receptors^30^. This network and behavioral effect of interneuronal α1A signaling in the BLA, in complement with earlier work implicating α1 signaling and LC-derived noradrenaline in valence assignment, suggests BLA interneurons may serve to govern preferential engagement of valence-specific BLA ensembles by modulating the local oscillatory network states. Here, we combine targeted cell-specific manipulations and closed-loop optogenetics with *in vivo/ex vivo* electrophysiology and behavior to demonstrate the crucial contribution of PV interneurons in the BLA to local computations governing the directionality of valence assignment. These data demonstrate a novel cellular mechanism mediating the ability of neuromodulators to govern valence assignment through a local BLA process involving specific interneuron-driven oscillatory states.

## Results

### LC-BLA noradrenaline drives real-time negative valence assignment

To address whether BLA α1A receptors are responsible for LC activation-induced real-time negative valence assignment^26^, we stereotaxically injected an adeno-associated viral vector guiding Cre-dependent ChrimsonR-tdTomato (or EYFP control) virus unilaterally into the LC of mice expressing Cre under the dopamine beta hydroxylase (Dbh) promoter. Additionally, a combined optic fiber and local field potential (LFP) recording electrode (optrode) was stereotaxically implanted in the BLA for LC-BLA noradrenergic terminal (LC-BLA^NE^) activation with simultaneous LFP recording (Figure 1A-C). To investigate the effect of LC-BLA^NE^ on real-time valence assignment, we employed a modified two-bottle choice task which 1) provides real-time assessment of affective information and 2) can disambiguate valence from salience. Following recovery, mice were placed in a two-bottle choice task (Figure 1B), during which they were presented with two identical water bottles to assess natural side preference^47^. Both bottles were fitted with infrared sensors surrounding pressure-sensitive lick spouts for millisecond-resolution recording of lick activity. Following 24 hours, licks from the first day’s preferred bottle (determined by bout numbers) were paired with a 3 second, 10 Hz, 60% pulse width (640nm, 5mW) optogenetic stimulation of LC-BLA^NE^ terminals.

**Figure 1.**
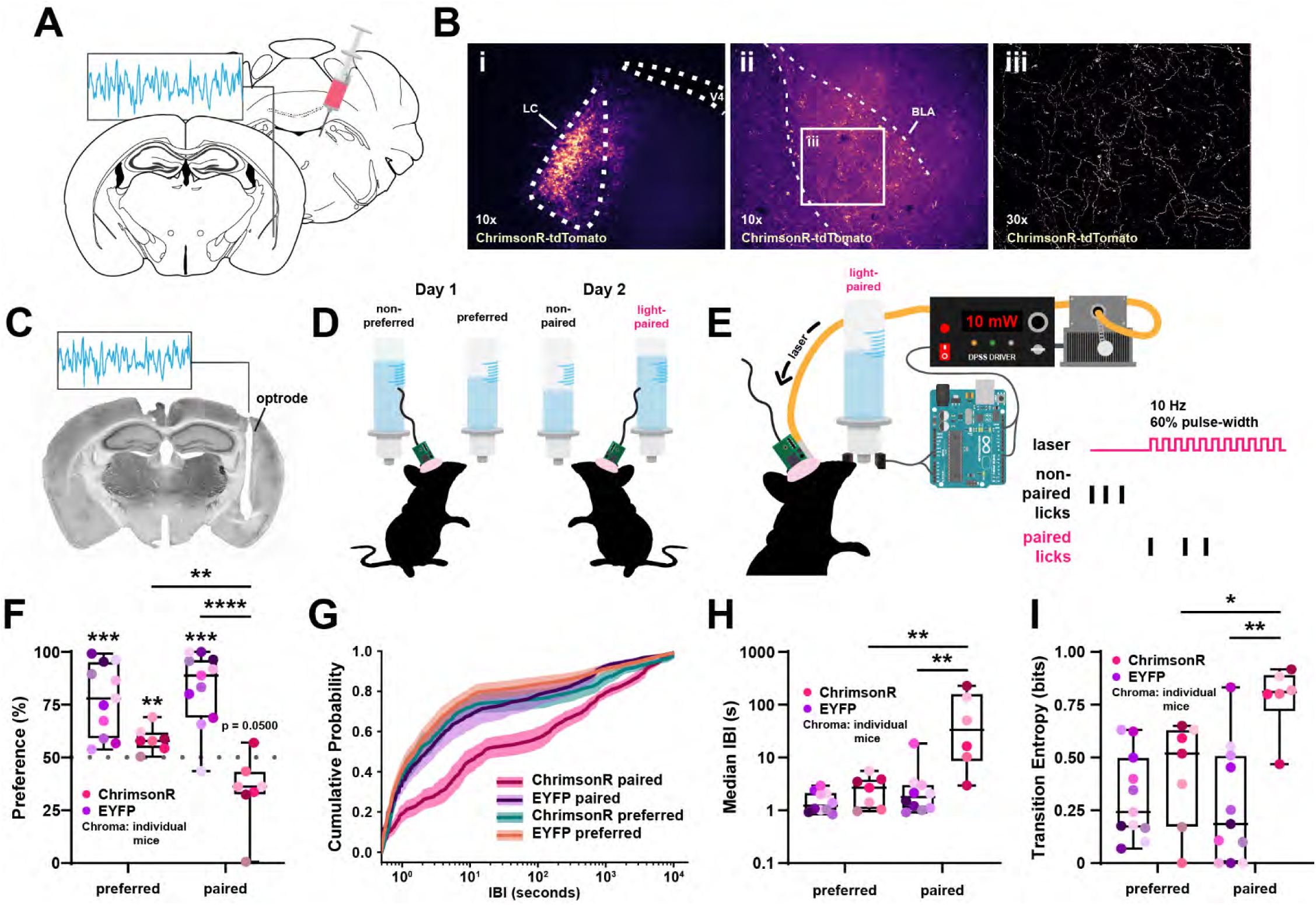
LC-BLA^NE^ facilitates negative valence assignment. **A.** Schematic illustrating viral injection and optrode implantation strategy. **B.** Representative images showing targeted expression of ChrimsonR-tdT in the LC (i) and its projections to the BLA (ii). iii. Magnified view of the boxed region in ii, highlighting dense LC terminal ChrimsonR-tdT expression in the BLA. **C.** Representative coronal section showing optrode placement in the BLA. **D,E.** Schematic of the experimental design in which tethered animals freely access two identical water sources during continuous electrophysiological recording across 48 hours. Licks from day 1’s preferred bottle are paired with optogenetic stimulation of LC-BLA terminals on day 2. **F.** Quantification of bottle preference as the percentage of total bouts across bottles per day and condition. **G.** Cumulative distribution of inter-bout intervals (IBI, seconds) across bottles and conditions. Lines = mean, shaded regions = SEM. **H.** Median IBI values (seconds) across bottles and conditions. **I.** Quantification of Shannon entropy (bits) for bottle transitions across bottles and conditions. For all box plots, hue = condition; chroma = individual animals. *, **, ***, **** = p < 0.05, 0.01, 0.001, 0.0001.

Animals displayed a natural preference for one bottle over the other during the first 24 hours that was significantly different from chance (Figure 1D, Statistics Table 1). However, during the second 24 hours, mice that were subjected to LC-BLA^NE^ terminal photostimulation during licks from the previously preferred bottle showed a reversal of bottle preference, as the percentage of lick bouts from the light-paired bottle was reduced to below chance (Figure 1D, Statistics Table 1), suggesting a real-time aversion to the light-paired bottle. In contrast, control animals expressing EYFP instead of ChrimsonR that received light delivery during licks from their previously preferred bottle maintained their initial bottle preference. Additionally, LC-BLA^NE^ terminal activation increased median inter-bout-interval (IBI) durations, suggesting an aversion to sustained engagement with the light-paired bottle (Figure 1E,F, Statistics Table 1). To assess whether this reflected a broader restructuring of licking behavior, we compared the full IBI distributions at the paired and preferred bottles using the two-sample Kolmogorov-Smirnov (KS) test. Mice exposed to LC-BLA^NE^ terminal activation showed consistently large per-mouse KS distances, indicating a robust shift in IBI structure at the light-paired bottle (Statistics Table 1). EYFP controls exhibited smaller but statistically significant KS distances (Statistics Table 1), which likely reflected minor variability across days rather than a meaningful update in licking structure.

To rule out the possibility that reduced engagement with the light-paired bottle was due to off-target effects such as motor suppression or satiety, we quantified the Shannon entropy of bottle transitions. Specifically, we asked: once a mouse resumes licking, how predictable are its choices? If stimulation truly drives negative valence assignment, we reasoned it should increase entropy by breaking the mouse’s structured preference for the previously preferred bottle, resulting in more random, less preferential transitions. In contrast, if stimulation merely induces satiety or motor suppression, it might temporarily reduce licking but should not change the underlying pattern of the preference, thus leaving entropy unchanged. As expected, animals from both conditions on day 1 showed highly structured and predictable bottle transitioning aligned with a developing preference, reflected in low Shannon entropy values (Figure 1G). This indicates that mice maintain a consistent bottle engagement strategy rather than exploring randomly, consistent with a goal-directed engagement. Further, EYFP control animals exposed to light delivery during licks from the previously preferred bottle on day 2 maintained a similar bottle engagement strategy, with transition entropy levels comparable to day 1, reflecting preservation of structured, preference-driven behavior. In contrast, animals exposed to LC-BLA^NE^ terminal stimulation displayed markedly increased transition entropy at the light-paired bottle (Figure 1G, Statistics Table 1). This shift suggests LC-BLA^NE^ stimulation elicited an aversive association with the previously preferred bottle, manifesting as a breakdown in structured, preference-driven engagement, rather than a general reduction in consummatory behavior.

Finally, to dissociate real-time negative valence from conditioned aversion and salience coding, we analyzed IBIs and bout durations using Bayesian hierarchical models. For IBIs, we fit a Bayesian Gaussian mixed-effects model on log-transformed values with random intercepts and slopes for each animal, allowing us to account for between-subject variability. This analysis revealed no credible effect of bout progression (Supplemental Figure 1A, Statistics Table 1), indicating that prolonged pauses between bouts emerged immediately rather than accumulating gradually across repeated stimulations. Parallel analyses of log-transformed bout durations likewise showed no consistent effect of within-session bout progression (Supplemental Figure 1B, Statistics Table 1), though did reveal a reduction in bout durations between days in both conditions (Supplemental Figure 1C, Statistics Table 1), suggesting that the vigor of bottle engagement did not systematically diminish over repeated trials. Together, these analyses support the conclusion that LC-BLA^NE^ terminal photostimulation drives an immediate, real-time negative valence signal, dissociable from conditioned aversion, salience coding, or nonspecific reductions in consummatory behavior.

**Supplemental Figure 1.**
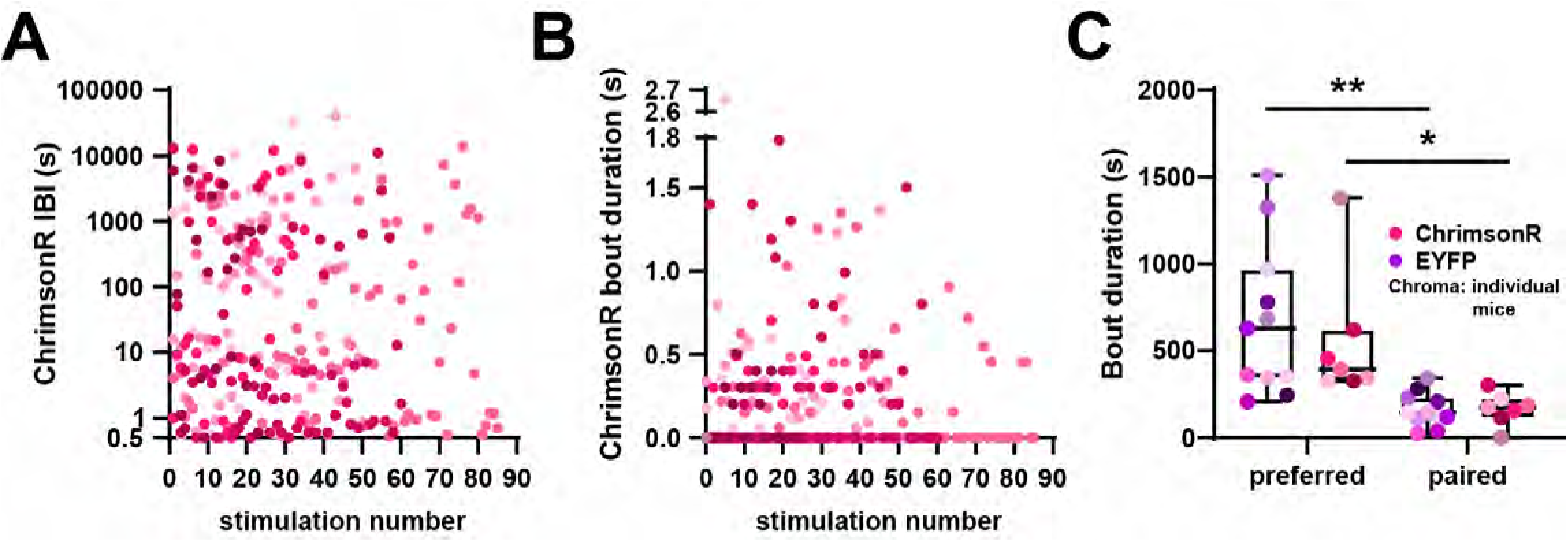
IBI and bout duration do not scale with successive stimulations. **A,B.** IBI (A) and bout durations (B) plotted as a function of stimulation number. Bout durations equal to 0 indicate single-lick bouts. **C.** Quantification of mean bout durations across bottles per day and condition. Hue = condition, chroma = individual animals. *, ** = p < 0.05, 0.01.

### LC-derived norepinephrine drives negative valence assignment through α1A receptors on BLA interneurons

Earlier work by McCall et al. (2015) demonstrates that tonic photostimulation of LC noradrenergic cell bodies is sufficient to produce real-time place aversion^26^. Moreover, systemic administration of the non-selective α1 receptor antagonist, prazosin, but not the broad β adrenergic receptor antagonist, propranolol, blocks this effect. Given our previous work showing BLA PV α1A receptors’ involvement in fear-associated behavior^30^, we asked here whether BLA α1A receptor signaling underlies LC-derived norepinephrine’s ability to facilitate real-time negative valence assignment. To investigate whether LC-BLA^NE^ drives negative valence assignment *via* BLA interneuronal α1A signaling, we crossed Dbh-Cre mice with *ADRA1A* knockout (*ADRA1A^-/-^*) mice^48^ to generate Dbh-Cre mice lacking α1A receptors (Dbh-Cre^α1AKO^), including in BLA PV interneurons (Figure 2A, Supplemental Figure 2). Note that we showed previously that α1A receptor expression is specific to GABA interneurons^30^. We drove expression of a Cre-dependent ChrimsonR-tdTomato with AAV-ChrimsonR-tdTomato injection in the LC of Dbh-Cre^α1AKO^ mice and tested whether LC-BLA^NE^ terminal stimulation in the α1A knockout mouse facilitates negative valence assignment in the two-bottle choice task. Dbh-Cre^α1AKO^ mice displayed a similar significant preference for one bottle over the other in the absence of photostimulation on day 1 (Figure 2B, Statistics Table 1). Unlike the Dbh-Cre mice with wild-type α1A receptor expression, LC-BLA^NE^ terminal photostimulation during licks from the previously preferred bottle on day 2 failed to reverse the bottle preference in the Dbh-Cre^α1AKO^ mice, (Figure 2B, Statistics Table 1). Assessment of full IBI distributions and medians illustrated no significant differences between Dbh-Cre^α1AKO^ and EYFP mice (Figure 2C,D, Statistics Table 1). Moreover, Dbh-Cre^α1AKO^ mice maintained similarly strong Shannon entropy across days, despite LC-BLA^NE^ terminal photostimulation (Figure 2E, Statistics Table 1). Taken together, these data suggest that LC-BLA^NE^ terminal photostimulation drives real-time negative valence assignment *via* BLA interneuronal α1A receptors, with PV interneurons representing a likely major cellular contributor given their demonstrated α1A expression and their known involvement in α1A-mediated negatively valenced behaviors.

**Figure 2.**
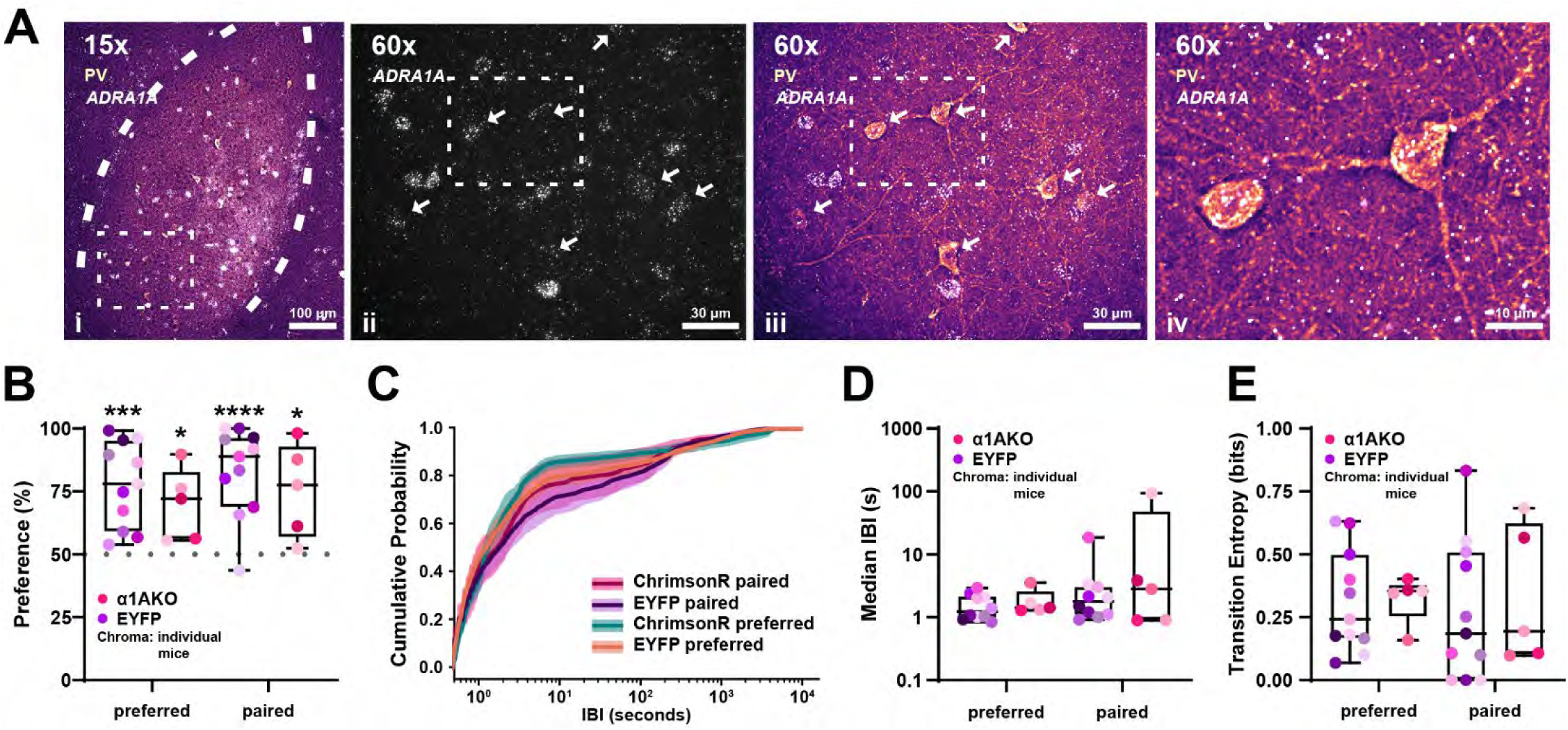
LC-BLA norepinephrine facilitates negative valence assignment *via* interneuronal α1A receptors. **A.** Representative dual Fluorescent In Situ Hybridization (FISH) and immunofluorescence images confirm *ADRA1A* transcripts are expressed in PV interneurons. (i) Low-magnification (15x) overview. (ii, iii) High-magnification (60x) images from the inset in (i), depicting *ADRA1A* transcripts alone (ii) and merged with PV immunolabeling (iii). (iv) Zoomed-in view of inset in ii/iii. **B.** Quantification of bottle preference as the percentage of total bouts across bottles per day and condition. **C.** Cumulative distribution of IBIs illustrating means (lines) and standard errors (shaded regions) across bottles and conditions. **D.** Median IBI values (seconds) across bottles and conditions. **E.** Quantification of Shannon entropy (bits) for bottle transitions across bottles and conditions. For all box plots, hue = condition; chroma = individual animals. *, **, ***, **** = p < 0.05, 0.01, 0.001, 0.0001.

**Supplemental Figure 2.**
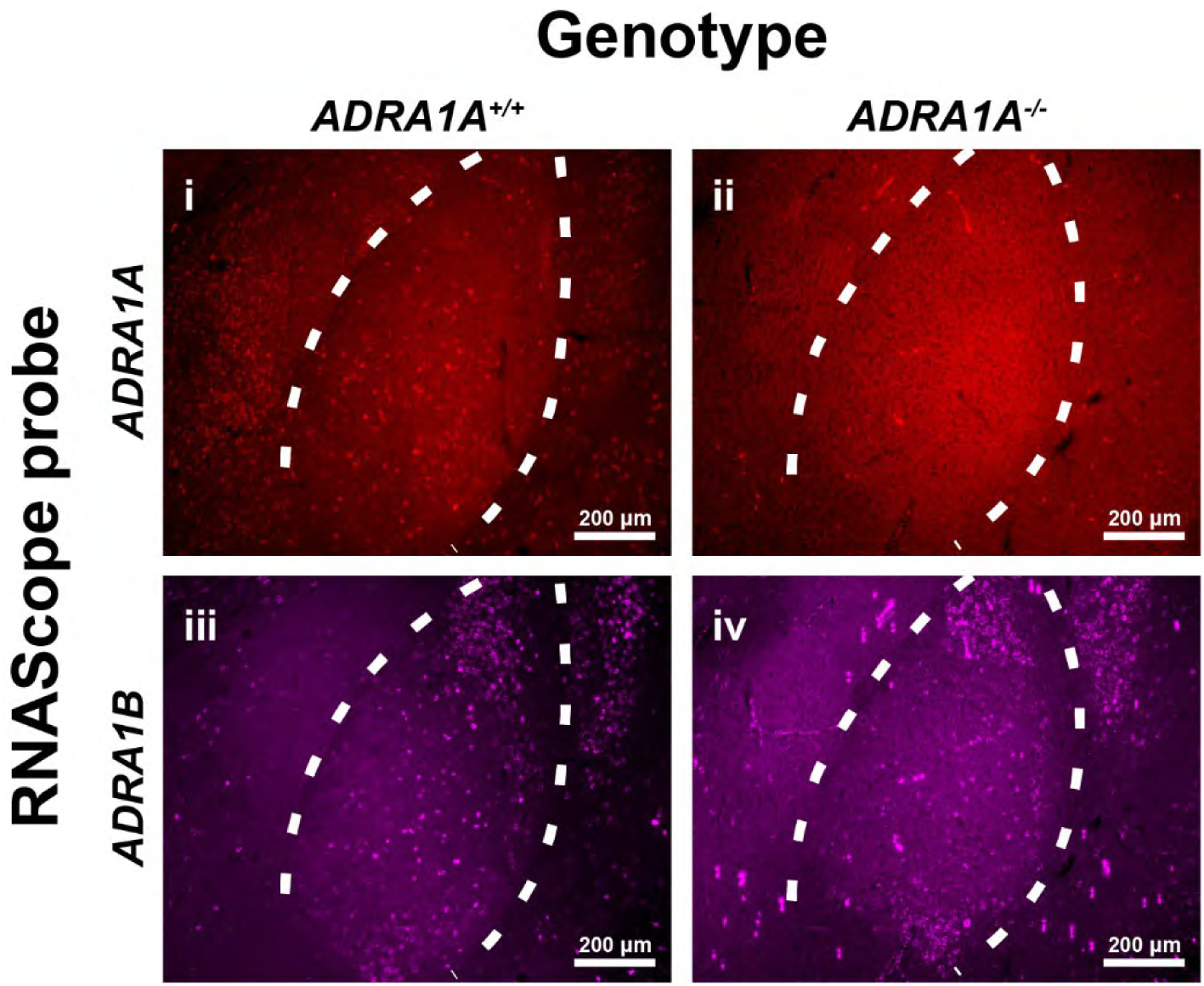
α1 adrenergic receptor expression in *ADRA1A*^+/+^ and *ADRA1A^-/-^*mice. Representative 10x magnification FISH images illustrating differential α1A and α1B adrenergic receptor subtype mRNA in the BLA of *ADRA1A^+/+^* (wild-type) and *ADRA1A^-/-^* (knockout) mice. (i, ii) *ADRA1A* mRNA in *ADRA1A* wild-type (i) and knockout (ii) mice. (iii, iv) *ADRA1B* mRNA in *ADRA1A* wild-type (iii) and knockout (iv) mice. Scale bars indicate 200 μm. Dashed lines demarcate the boundary of the BLA.

### LC-derived norepinephrine modulates BLA oscillatory network states

We recently showed that Gq signaling downstream of α1A receptor activation following intra-BLA norepinephrine administration and BLA PV interneuron-specific hM3D activation suppresses local “fast gamma” (70-120 Hz) oscillatory power *in vivo*^30^. Given the extensive evidence that BLA network states are associated with distinct behavioral states, including those involved in valence processing^30,32,36,37,39–41,51^, we asked whether LC-BLA^NE^ terminal photostimulation may influence BLA network states associated with valence assignment in the two-bottle choice task.

Consistent with our earlier observations, BLA LFP recordings in animals that received LC-BLA^NE^ terminal stimulation exhibited suppression of local “fast gamma” oscillations (70-120 Hz) for 1-to-5 seconds post-lick, outlasting the 3-second stimulation period. (Figure 3A-C, Statistics Table 2). This suppression closely parallels our previous findings demonstrating that BLA PV interneurons mediate Gq-driven suppression of fast gamma oscillations following norepinephrine application or chemogenetic activation^30^. In contrast, light delivery in Dbh-Cre mice expressing control EYFP failed to suppress fast gamma oscillatory power (Figure 3A-C, Statistics Table 2).

**Figure 3.**
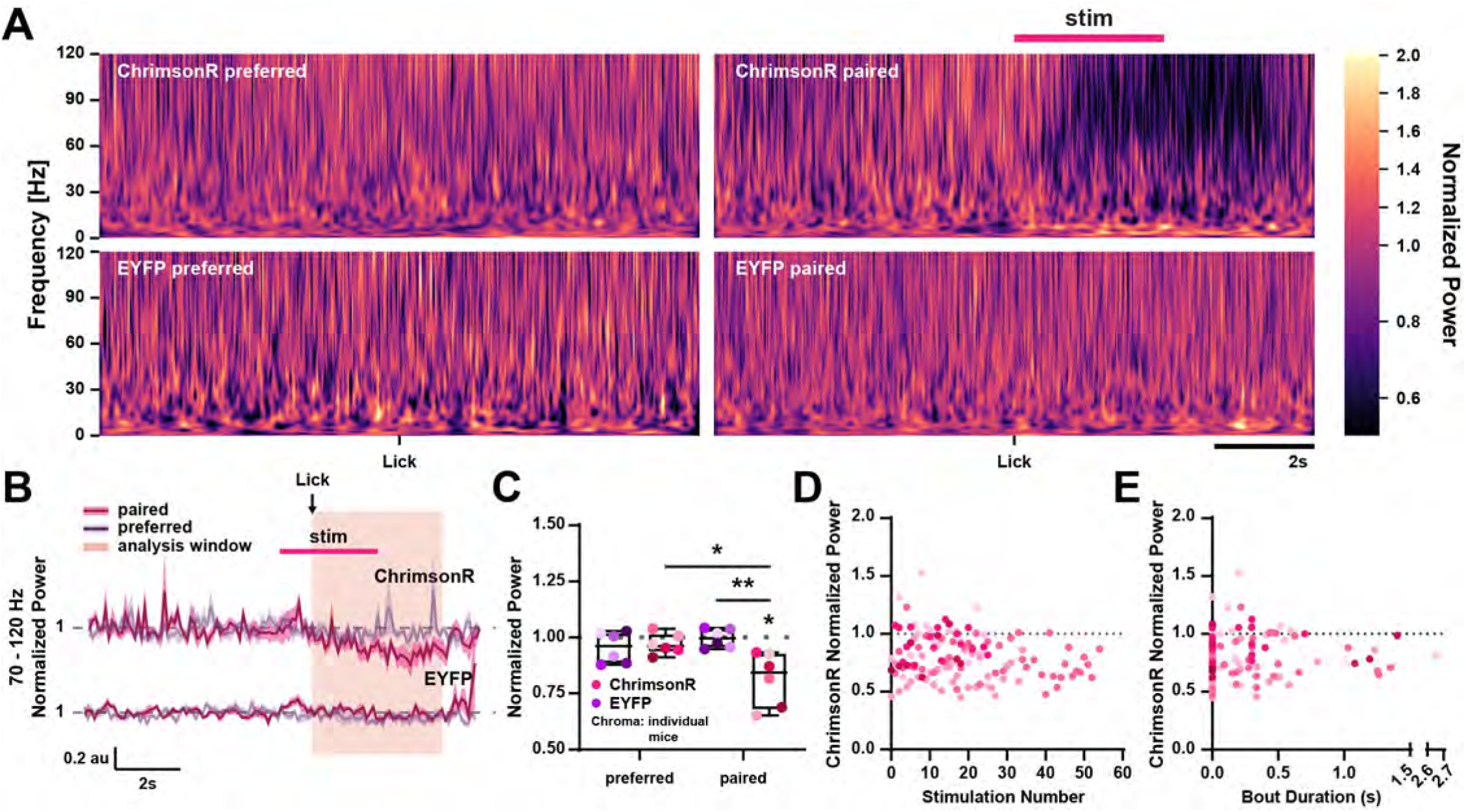
LC-BLA norepinephrine suppresses BLA fast gamma oscillatory power. **A.** Average peri-lick spectrograms from a single representative mouse per condition initiating new lick bouts at the previously preferred (left) and light-paired (right) bottles. **B.** Average baseline-normalized 70-120 Hz peri-lick power time series plots across animals, shown per bottle and condition (line = mean; shaded region = SEM). Values are normalized to the 3-second pre-lick baseline. Analysis window = 1-5 seconds post-lick. **C.** Quantification of normalized 70-120 Hz power within shaded analysis window in B. **D,E.** Per-bout 70-120 Hz power in ChrimsonR-expressing mice plotted as a function of stimulation number (D) or bout duration (E) per animal. For box and scatter plots, hue = condition; chroma = individual animals. *, **, = p < 0.05, 0.01.

To confirm that light-evoked suppression of BLA fast gamma rhythms reflects a real-time rather than conditioned process, we tested whether successive light deliveries resulted in greater suppression magnitude across animals (Figure 3D). We fit a Bayesian hierarchical linear model predicting normalized 70-120 Hz power (centered on baseline = 1) from trial (lick bout) progression with varying intercepts by mouse to account for between-subject variability, while slopes were constrained to be equal across animals to improve convergence given near-zero slope variance. The model revealed no consistent change in gamma suppression across trials: the fixed effect of bout number was nearly zero, providing no strong evidence that suppression increased or decreased with repeated stimulation (Figure 3D; Statistics Table 2). Random effects indicated moderate variability in intercepts across mice – perhaps due to variability in viral or optical targeting (Statistics Table 2). These findings indicate that while optogenetic stimulation reliably induces fast gamma suppression, the magnitude of suppression remains stable across repeated trials, supporting the interpretation that the suppression is real-time and stimulus-locked, rather than a progressively learned or accumulating effect.

Finally, to disambiguate whether light-evoked suppression of BLA oscillatory power primarily signals valence or stimulus salience, we tested whether oscillatory power predicted bottle engagement vigor. Specifically, we examined whether high gamma power during bouts from the light-paired bottle was associated with bout duration (Figure 3E). We fit a Bayesian hierarchical linear model predicting log-transformed bout duration from gamma power (centered at baseline = 1), with varying intercepts by mouse and a common slope estimated across animals (see Methods). The posterior distribution of the fixed effect indicated high uncertainty and overlaid with zero, suggesting no consistent relationship across animals (Statistics Table 2). These findings suggest that fast gamma power fluctuations were not predictive of bout duration, arguing against a salience-like role for light-evoked BLA oscillatory suppression.

Consistent with a role for interneuronal α1A signaling in shaping BLA oscillations, LC-BLA^NE^ terminal activation in animals lacking α1A receptors failed to suppress BLA fast gamma oscillatory power (Figure 4A-C, Statistics Table 2). Further, we fit a Bayesian hierarchical linear model predicting 70-120 Hz power (centered on baseline = 1) from trial progression. The model revealed only modest evidence for trial-wise suppression and the slope’s 95% credible interval included zero (Statistics Table 2), suggesting that fast gamma power remained largely stable across repeated bouts in the absence of α1A signaling.

**Figure 4.**
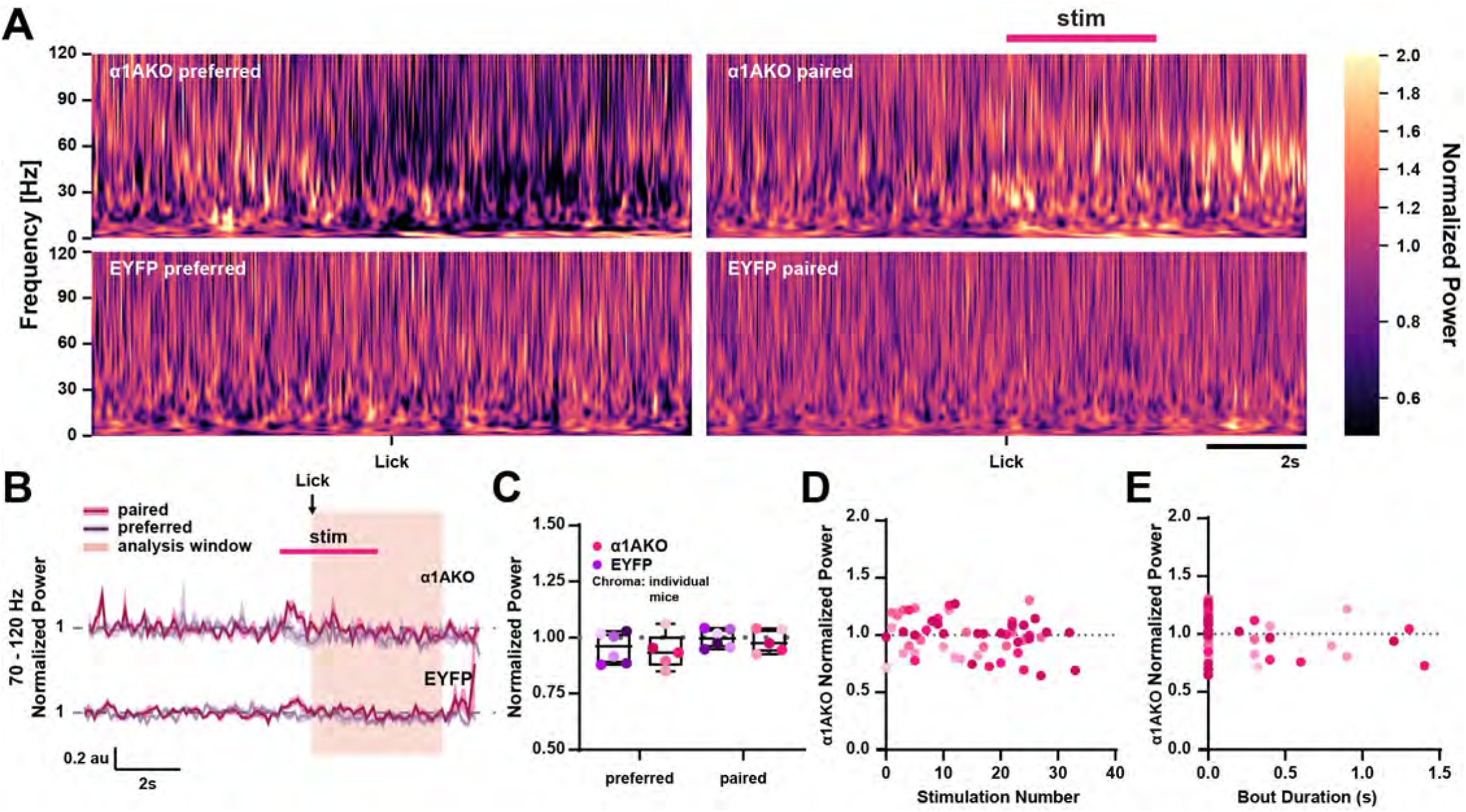
*ADRA1A* knockout blocks LC-BLA norepinephrine-driven fast gamma suppression. **A.** Average peri-lick spectrograms from a single representative mouse per condition initiating new lick bouts at the previously preferred (left) and light-paired (right) bottles. **B.** Average baseline-normalized 70-120 Hz peri-lick power time series plots across animals, shown per bottle and condition (line = mean, shaded region = SEM). Values are normalized to the 3-second pre-lick baseline. Analysis window = 1-5 seconds post-lick. **C.** Quantification of normalized 70-120 Hz power within shaded analysis window in B. **D,E.** Per-bout 70-120 Hz power in ChrimsonR-expressing *ADRA1A* knockout mice plotted as a function of stimulation number (D) or bout duration (E) per animal. For box and scatter plots, hue = condition; chroma = individual animals.

To test the association between fast gamma activity and behavior, we fit a separate Bayesian hierarchical model predicting log-transformed bout duration from fast gamma power (centered at baseline = 1), again allowing for varying intercepts by mouse. This model revealed no consistent relationship between fast gamma and bout duration (Figure 4E; Statistics Table 2), indicating that fast gamma power also fails to reliably predict behavioral output in α1A-deficient animals. Together, these data suggest that LC-BLA^NE^ suppresses 70-120 Hz oscillatory power *via* putative PV α1A signaling in a manner that reflects real-time negative valence assignment.

### BLA PV interneurons govern positive valence assignment and associated gamma rhythms

Thus far, our data suggest that BLA gamma rhythm suppression, likely driven by Gq signaling in local PV interneurons, underlies real-time negative valence assignment. This aligns with our previous work demonstrating that chemogenetic and optogenetic modulation of BLA PV interneuron dynamics can facilitate negatively valenced behaviors^30,32^. Notably, we also found that optogenetically driving BLA PV interneurons at different frequencies bidirectionally influences affective states^32,37,41^, raising the possibility that these interneurons may also contribute to computations supporting positive valence assignment. To test this hypothesis, we next examined whether targeted BLA PV interneuron knockdown (PVKD) influences bottle preference in our two-bottle choice task when presented with a positive stimulus (Figure 5A).

**Figure 5.**
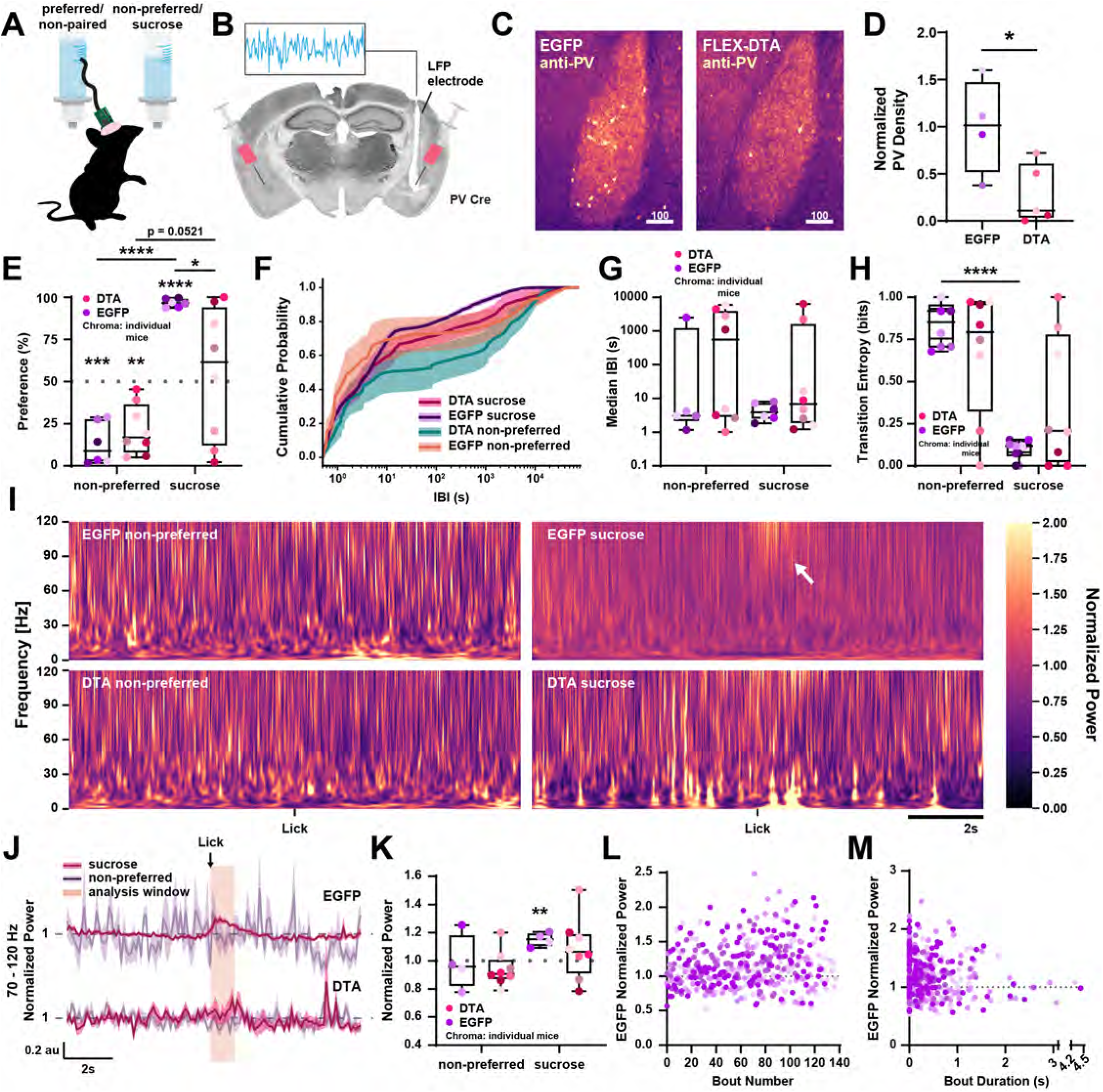
BLA PV-driven 70-120 Hz oscillatory power signals positive valence assignment. **A.** Schematic of experimental design in which tethered animals freely access two bottles (water/water followed by water/sucrose) during continuous electrophysiological recording. **B.** Representative coronal section and schematic illustrating bilateral BLA virus injection and unilateral recording electrode placement. **C.** Representative images showing PV immunolabeling in control (left) and PV knockdown (right) mice. Scale bar = 100 μm. **D.** Quantification of BLA PV density normalized to the average of control animals. **E.** Quantification of bottle preference as the percentage of total bouts across bottles per day and condition. **F.** Cumulative distribution of IBIs illustrating means (lines) and SEMs (shaded regions) across bottles and conditions. **G.** Median IBI values (seconds) across bottles and conditions. **H.** Quantification of Shannon entropy (bits) for bottle transitions across bottles and conditions. **I.** Average peri-lick spectrograms from a single representative mouse per condition initiating new lick bouts at the previously non-preferred (left) and sucrose-paired (right) bottles. White arrow denotes effect. **J.** Average baseline-normalized 70-120 Hz peri-lick power time series plots across animals, shown per bottle and condition (line = mean; shaded region = SEM). Values are normalized to the 3-second pre-lick baseline. Analysis window = 0-1 seconds post-lick. **K.** Quantification of normalized 70-120 Hz power in shaded analysis window in J. **L,M.** Per-bout 70-120 Hz power in control mice plotted as a function of bout number (L) or bout duration (M) per animal. For all box and scatter plots, hue = condition; chroma = individual animals. *, **, ***, **** = p < 0.05, 0.01, 0.001, 0.0001.

To selectively lesion BLA PV interneurons, we used a viral strategy for Cre-dependent expression of diphtheria toxin fragment A (DTA), the component of diphtheria toxin responsible for inhibiting protein synthesis and promoting cell death^52,53^. Adult PV-Cre mice received bilateral BLA injections of either AAV2/9-mCherry-FLEX-DTA or EGFP control virus, along with unilateral implantation of an LFP recording electrode to monitor rhythmic network activity. Quantification of PV cell survival showed that this Cre-dependent DTA strategy resulted in a 72.09 ± 14.21% reduction in PV cell counts compared to controls (Figure 5B-D, Statistics Table 3).

Following recovery from surgery, animals underwent the two-bottle choice task (as in the LC-BLA^NE^ experiments), during which they were presented with two identical water bottles to determine natural side preference for 24 hours. In contrast to the prior experiments, the contents of each animal’s day 1 non-preferred bottle were removed and replaced with a 20% sucrose solution. Again, animals with and without normal BLA PV interneuron counts displayed a natural preference for one bottle over the other during the first 24 hours that was significantly different from chance (Figure 5E, Statistics Table 3). Animals with wild-type BLA PV interneuron counts then demonstrated a marked reversal of bottle preference towards the sucrose-containing bottle during the second 24 hour period. However, PVKD animals failed to show the clear sucrose preference observed in controls, instead displaying a wide range of day 2 bottle preferences that were statistically indistinguishable from chance and significantly different from controls (Figure 5E, Statistics Table 3). Additionally, while group-level IBI cumulative distribution plots overlapped throughout the curve (Figure 5F), and no differences were appreciable through measures of central tendency, a Wilcoxon signed-rank test on per-mouse KS distances revealed statistically significant within-subject differences between the paired and non-preferred bottles (Statistics Table 3). However, given the subtle nature of these differences and the absence of consistent population-level shifts, the functional relevance of this statistical result is likely limited and may reflect heterogenous, individually idiosyncratic variation. In contrast, Shannon entropy analysis of bottle transitions revealed a marked reduction in EGFP mice, with entropy dropping from 0.866 ± 0.053 at the non-preferred bottle to 0.099 ± 0.022 upon introducing 20% sucrose, reflecting a near-deterministic pattern of repeated returns to the paired bottle, both through self-transitions and directed switches (Figure 5H, Statistics Table 3). In contrast, PVKD animals on average did not exhibit a comparable reduction in Shannon entropy at the sucrose-paired bottle, instead displaying highly variable entropy values that were not significantly different from those at the non-preferred bottle or the EGFP paired bottle (Figure 5H, Statistics Table 3).

Earlier, our laboratory reported selective BLA PV interneuron knockdown does not grossly alter baseline local network activity^31^. Consistently, BLA LFP activity recorded during licks from non-preferred water bottles on day 1 does not differ significantly from pre-lick baseline in either wild-type or PVKD animals (Figure 5I-K, Statistics Table 3). Moreover, no significant differences were observed between conditions. However, when exposed to sucrose, wild-type mice exhibit a roughly 13% increase in 70-120 Hz oscillatory power during the first 600ms post-lick across engagements, which differs significantly from pre-lick baseline. In contrast, PVKD animals fail to potentiate 70-120 Hz power during sucrose-paired licks (Figure 5I-K, Statistics Table 3). These data suggest that PV interneurons in the BLA may serve to coordinate valence-associated changes in network states rather than baseline activity.

To test whether the sucrose-evoked increase in 70-120 Hz power reflects a real-time versus conditioned response, we fit a Bayesian hierarchical model predicting normalized fast gamma power from bout progression in EGFP controls, with random intercepts for each animal to account for between-subject variability. We observed a modest but statistically significant increase in 70-120 Hz power across successive bouts, though the magnitude of this increase was very small (∼0.14% increase per bout) and variance explained was minimal (Bayes R^2^ = 0.04; Statistics Table 3), suggesting that this effect may be less biologically meaningful (Figure 5L). Overall, BLA fast gamma remained relatively stable across sucrose exposures, supporting a role for fast gamma in signaling stimulus positivity in real-time without clear evidence of learning-dependent scaling. To determine whether sucrose-driven enhancements in 70-120 Hz power reflect valence vs salience, we fit a Bayesian hierarchical model predicting log-transformed bout duration from fast gamma power (centered on baseline = 1) with random intercepts for each mouse. The posterior slope estimate was negative but highly uncertain (Statistics Table 3), indicating no reliable relationship between fast gamma and bout duration across animals (Figure 5M). Together, these findings suggest that enhanced 70-120 Hz BLA activity specifically reflects real-time positive valence assignment.

Mounting evidence demonstrates that local BLA rhythms and outputs contribute to motivational processes^41,54,55^. Therefore, it is not clear whether deletion of BLA PV interneurons resulted in impaired sucrose preference due to impairments in reward seeking behavior or valence assignment. To address this, we tested PVKD animals’ preference for 20% sucrose under food deprivation to increase appetitive motivation. We reasoned that if BLA PV deletion impairs appetitive motivation, then food-deprived animals should maintain a lack of preference for 20% sucrose over water. However, when reduced to 80-90% bodyweight, both wild-type and PVKD animals displayed a roughly 99% preference for 20% sucrose over water (Figure 6A, Statistics Table 3). Additionally, similar to sated control animals, neither food-deprived group displayed a significant shift in IBI central tendency, though Wilcoxon signed-rank tests on per-mouse KS distances revealed a statistically significant difference between paired and non-preferred bottle IBI distribution shapes in EGFP, but not PVKD mice - despite a visual divergence in cumulative distribution plots within the top 20th percentile (Figure 6B,C, Statistics Table 3). To further characterize the extent to which motivational drive under food deprivation reinstated directed, preference driven behavior, we analyzed bottle transition entropy. Both EGFP controls and PVKD mice exhibited highly structured transitions under food deprivation at the sucrose-paired bottle, reflected in lower Shannon entropy values compared to the non-preferred bottle (Figure 6D, Statistics Table 3). These data support the idea that PV interneurons in the BLA contribute to valence assignment rather than reward seeking related to motivated behaviors.

**Figure 6.**
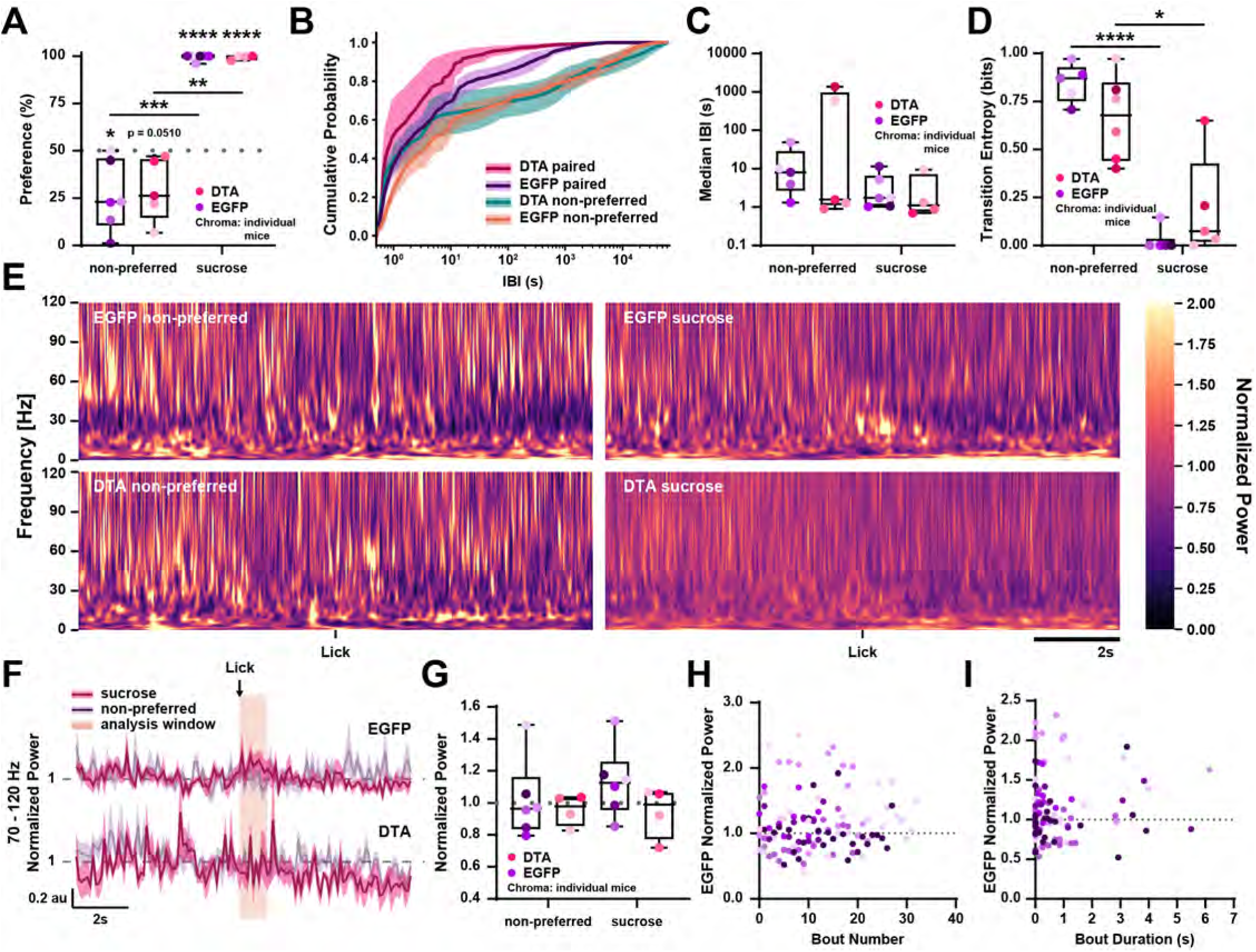
Increased motivation restores food-seeking behavior without concomitant 70-120 Hz power enhancement. **A.** Quantification of bottle preference as a percentage of total bouts across bottles per day and condition. **B.** Cumulative distribution of IBIs illustrating means (lines) and SEMs (shaded regions) across bottles and conditions. **C.** Median IBI values (seconds) across bottles and conditions. **D.** Quantification of Shannon entropy (bits) for bottle transitions across bottles and conditions. **E.** Average peri-lick spectrograms from a single representative mouse per condition initiating new lick bouts at the previously non-preferred (left) and sucrose-paired (right) bottles. **F.** Average baseline-normalized 70-120 Hz peri-lick power time series across animals, shown per bottle and condition (line = mean, shaded region = SEM). Values are normalized to the 3-second pre-lick baseline. Analysis window = 0-1 seconds post-lick. **G.** Quantification of normalized 70-120 Hz power in shaded analysis window in F. **H,I.** Per-bout 70-120 Hz power in control mice plotted as a function of bout number (H) or bout duration (I) per animal. For all box and scatter plots, hue = condition; chroma = individual animals. *, **, ***, **** = p < 0.05, 0.01, 0.001, 0.0001.

Given that reward-seeking behavior was restored by caloric deficit, we asked whether the fast network dynamics observed during sucrose-paired licks in sated controls would similarly re-emerge in fasted PVKD animals. We reasoned that if 70-120 Hz oscillatory power selectively reflects valence-associated BLA computations, then enhancing motivational drive should not be sufficient to restore this rhythm. As in our sated animals, BLA LFP activity during licks from the non-preferred bottle on day 1 did not differ significantly from either the pre-lick baseline or between groups in fasted PVKD animals (Figure 6E-G, Statistics Table 3). Moreover, surprisingly, neither PVKD nor control animals exhibited a significant enhancement in fast gamma power during sucrose-paired licks, despite apparent reward-seeking behavior in both conditions (Figure 6E-G, Statistics Table 3). Consistent with this, Bayesian hierarchical models showed no systematic change in 70-120 Hz power across successive sucrose-paired licks and no reliable relationship between fast gamma power and log-transformed bout duration (Figure 6 H,I, Statistics Table 3). Together, these data indicate that 70-120 Hz power does not reflect motivation-associated BLA computations as caloric deprivation did not enhance fast gamma power, although they also reveal the unexpected finding that fasting abolishes the fast gamma enhancement normally seen during sucrose exposure in control animals.

Maintenance of food-seeking behavior in fasted controls despite the absence of fast gamma potentiation suggests that processes engaging motivated behavior may shift to need-based BLA computations from those supporting positive valence coding. In line with this interpretation, and consistent with our earlier characterization of BLA PV-driven beta (15-30 Hz) oscillatory activity as a motivational value-linked rhythm sufficient to drive reward-seeking behavior^41^, we observed enhanced beta oscillatory power during sucrose exposure only when it was combined with caloric deprivation (Supplemental Figure 3 A-B, E-F, Statistics Table 3). Specifically, fasted animals displayed on average a ∼20% increase in BLA beta power during licks from the sucrose-paired bottle, in contrast to licks from the non-paired bottle or from either bottle in the sated condition. Notably, Bayesian hierarchical modeling revealed beta power did not change across successive bouts (Supplemental Figure 3 C,G, Statistics Table 3) and showed no relationship with log-transformed bout duration in either the sated or fasted state (Supplemental Figure 3 D,H, Statistics Table 3), suggesting that this rhythm reflects a motivational state driven by sustained enhanced caloric need across the session, rather than incremental reward learning within the session. This pattern points to a functional reallocation of PV-driven BLA rhythms, in which fast gamma activity normally associated with positive valence is shifted toward beta oscillations that encode motivational value under caloric deprivation. Interestingly, this increase in beta power required BLA PV interneurons (Supplemental Figure 3 E,F), reinforcing the circuit mechanism identified in our previous work^41^ and questioning the necessity of BLA PV-driven beta rhythms for motivated action and learning given the maintenance of reward-seeking behavior in PVKD animals. Collectively, these findings suggest that while BLA PVKD disrupts sucrose preference, core motivational processes remain preserved, such that the caloric demands of fasting override the loss of positive affective valuation and sustain engagement with energy-rich food which does not involve a concomitant fast gamma oscillatory signature.

**Supplemental Figure 3.**
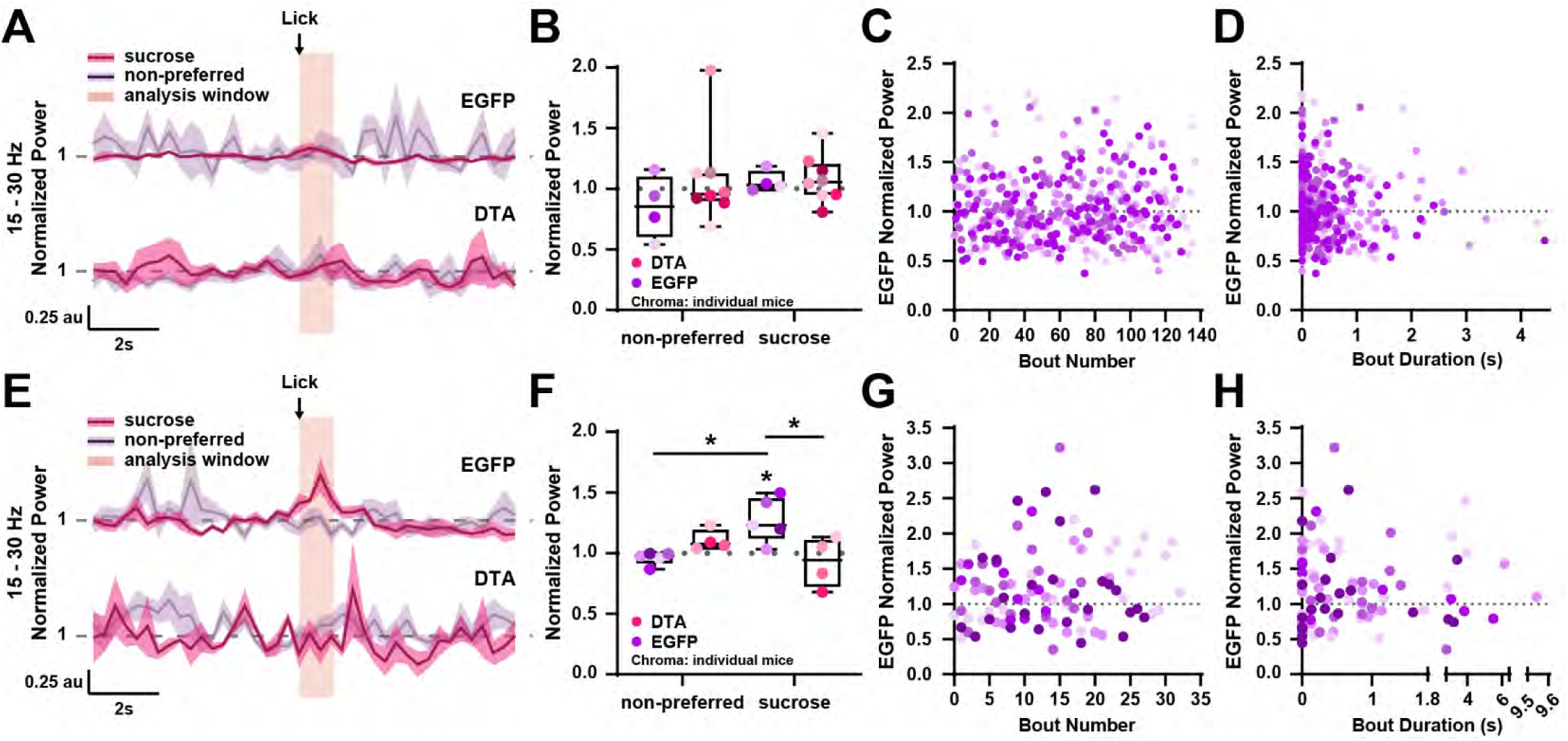
Increased motivation under caloric deprivation enhances BLA PV-driven beta oscillatory activity during sucrose consumption. **A,E** Average baseline-normalized beta (15-30 Hz) peri-lick power time series plots across animals, shown per bottle and viral condition in sated (A) and deprived (E) states (line = mean, shaded region = SEM). Values are normalized to the 3-second pre-lick baseline. Analysis window = 0-1 seconds post-lick. **B,F.** Quantification of normalized 15-30 Hz power within shaded analysis window in A,E. **C-D,G-H.** Per-bout 15-30 Hz power in control mice plotted as a function of bout number (C,G) or bout duration (D,H) per animal in sated (C-D) and deprived (G-H) states. For all box and scatter plots, hue = condition; chroma = individual animals. * = p < 0.05

## Discussion

Here, we identify a previously unrecognized role for neuromodulation of BLA inhibitory interneurons, particularly PV interneurons, in governing real-time valence assignment. Further, we demonstrate that this mechanism is capable of influencing both positive and negative valence assignment. Using a combination of closed-loop optogenetics, behavior, *in vivo* and *ex vivo* electrophysiology, and pharmacological manipulations, we show that LC-derived norepinephrine engages BLA interneuronal α1A receptors to drive real-time negative valence assignment and associated local fast gamma power suppression. In parallel, we demonstrate that BLA PV interneurons are also necessary for real-time positive valence assignment, as their selective ablation disrupts sucrose preference and the associated enhancement of local fast gamma power, despite preserved reward-seeking behavior under conditions of increased motivational drive. Together, these findings reveal that BLA PV interneurons dynamically regulate affective valuation in both directions, a process involving the orchestration of unique local network rhythms.

These findings challenge prevailing models of BLA valence coding that place glutamatergic BLA principal neurons at the center of valence processing^12,13,16–18,20–22^. Instead, our data reveal that BLA inhibitory neurons – which, for the most part, do not send long-range projections^49,50^ – critically contribute to valence assignment, despite prior comprehensive mapping of BLA valence-coding outputs failing to detect inhibitory populations^16,17,21^. This divergence highlights an underappreciated computational role of local BLA inhibitory circuits in valence processing and motivates a reevaluation of how network dynamics within the BLA contribute to valence assignment.

We propose an updated model of BLA valence computation, whereby local inhibitory microcircuits play a fundamental and previously underappreciated role in shaping valence assignment in real-time. We posit that the collective activity among local inhibitory interneurons, particularly PV interneurons, shapes BLA oscillatory dynamics in real-time to influence valence assignment *via* rhythm-dependent differential gain control of valence-specific BLA principal neurons. This means the same PV cell network, by shifting its rhythmic pattern, selectively biases the excitability of one principal neuron assembly over a competitor, thereby enabling rapid and flexible information routing in response to competing amygdalar inputs, enabling flexible navigation of dynamic environments where temporally convergent but opposing stimuli introduce ambiguity. In fact, this interpretation of BLA gamma function in valence assignment largely aligns with Feng and colleagues’ biophysical model of BLA gamma in which competing principal neuron assemblies are recruited by the local gamma oscillation^64^. This alignment further parallels empirical findings from other groups, offering a mechanistic link between dynamic gamma oscillatory activity and the BLA’s “valence tracking” property, where individual neurons and connected microcircuits respond differentially to stimuli with opposing valences^12,18,65^. This novel mechanism governing valence assignment at the level of BLA network states would confer real-time dynamic neural computations.

Similar gamma rhythmic BLA oscillatory network dynamics have been observed in fear states. In particular, we and others have shown that fear expression coincides with similar degrees of fast gamma suppression^30,39^, and that BLA PV Gq signaling is sufficient to drive this effect^30^. Building on this, studies of fear expression reveal that fast gamma suppression is accompanied by enhanced low theta power^32,36,37^ and theta-gamma coupling^39^ in the BLA. These findings suggest that fear-associated gamma suppression operates not in isolation but as part of a layered rhythm, one that nests within local theta. Our data extend this view by showing that fast gamma suppression arises more fundamentally during negative valence assignment, a computation that underlies fear. In this framework, fast gamma suppression provides a local marker of valence within the BLA, and when aligned with local theta, these components integrate into the circuit-level dynamics that give rise to the emergent state of fear.

The finding that BLA PV interneurons govern both positive and negative valence assignment raises important questions about the molecular mechanisms through which PV interneuron signaling shapes local network states and their capacity to bias valence computations. Previous data from our laboratory demonstrated that other neuromodulatory systems, including the dopaminergic system, are capable of modulating BLA network states^51^, representing complementary regulatory mechanisms. We demonstrate additional complexity mediating the effects of norepinephrine on BLA network states, involving Gq-coupled α1A adrenergic receptor signaling in local PV interneurons.

Here, we provide strong support for a model in which BLA PV interneuron activity dynamically regulates local network states to bidirectionally drive real-time valence assignment. Future studies leveraging cell-type-specific receptor deletions, *in vivo* imaging of neural activity and Ca^2+^ dynamics, and comprehensive pharmacological validation will be valuable to refine our mechanistic understanding further.

## Methods

### Animals

C57BL6/J (C57; Jax, Strain # 000664), B6.129P2-Pvalbtm1(cre)Arbr/J (PV-cre; The Jackson Laboratory, Strain # 017320), B6.FVB(Cg)-Tg(Dbh-cre)KH212Gsat/Mmucd (Dopamine beta hydroxylase (Dbh)-cre, MMRRC Stock # 036778-UCD), B6.129X1-Adra1atm1Pcs/J (ADRA1A^-/-^ (knockout); The Jackson Laboratory, Strain # 005039), Dbh-Cre^α1AKO^ (Dbh-Cre x *ADRA1A^-/-^* cross; generated in-house) mice used in this study were housed in an AAALAC-approved housing facility at Tufts University School of Medicine under a standard 12-hour light cycle with food and water provided *ad libitum*, unless specified otherwise. All procedures were approved by Tufts University’s Institutional Animal Care and Use Committee (IACUC) and all animals were regularly monitored by members of the Maguire Lab and Tufts Comparative Medicine Services animal care staff.

### Surgery

Mice at least 8-weeks-old were anesthetized by intraperitoneal injection of 100mg/kg ketamine + 10mg/kg Xylazine until unresponsive to foot pinch prior to placement in a mouse stereotaxic frame. The skull was exposed along the midline under aseptic conditions and craniotomies were performed over the BLA and/or LC for virus injection or electrode/optrode implantation.

#### Virus Injections

A pulled glass pipette with a 20 μm diameter opening carrying virus solution was lowered to the desired target (LC: AP - 5.22, ML ± 0.93, DV - 2.95 relative to dura; BLA: AP - 1.35, ML ± 3.3, DV - 5.1 relative to the skull at Bregma) at an approximate rate of 0.1 mm/second. Virus solution was manually pressure-injected at 100 nL/minute using a 10 cc syringe and flexible PTFE tubing. Immediately following injection, the virus pipette was allowed to rest for an additional 4 minutes before removing from the brain at an approximate rate of 0.1 mm/second.

#### Optrode Implantation

A combined LFP recording electrode (PFA-coated stainless steel wire (A-M Systems #792100) and pre-fabricated electrode interface boards (Pinnacle Technology cat. # 8201)) and 200 μm core, 0.39 NA implantable optic fiber terminating 1 mm above the recording site (ThorLabs cat. # CMXB10) or Neuronexus microelectrode array (A1×32-Poly2-5mm-50s-177-OZ32LP) with an integrated optic fiber cannula (50 μm core, 0.22 NA) terminating ∼ 825 μm above the bottom of the silicone shank (both referred to here as “optrode”) was implanted in/above the BLA (AP - 1.35, ML ± 3.40, DV - 5.1 (Pinnacle) or - 5.15 (Neuronexus) relative to skull at Bregma). Following injection and implantation, all hardware was affixed to the skull with dental cement A-M Systems, #525000 and #52600) before removal from the stereotaxic frame. Finally, animals were placed in a heated recovery chamber until conscious before returning to their home cages.

### Viral Vectors

Adeno-associated viral (AAV) vectors were obtained from Addgene and stored at −80 °C until use. For LC-BLA^NE^ experiments, AAV5-Syn-FLEX-rc[ChrimsonR-tdTomato] (Addgene, Cat. # 62723-AAV5, titer = 8.5 x 10^12^ GC/ml) or AAV1-Ef1a-DIO-EYFP (Addgene, Cat. # 27056-AAV1, titer = 2.5 x 10^13^ GC/ml) were injected unilaterally into the LC of Dbh-Cre and Dbh-Cre^α1AKO^ mice for optical control over LC terminals in the BLA. For PV knockdown experiments, AAV2/9-mCherry-FLEX-DTA (Addgene, plasmid only, Cat. # 58536, titer = 8.72 x 10^13^ GC/ml) or AAV9-hSyn-DIO-EGFP (Addgene, Cat. # 50465-AAV9, titer = 1.9 x 10^13^ GC/ml) were injected bilaterally into the BLA of PV-Cre mice for BLA PV neuron-specific knockdown.

### Electrophysiology

#### In vivo

Local field potential recordings were performed in awake, freely behaving mice using either a standard recording electrode (A-M Systems/Pinnacle) or a silicone microelectrode array (Neuronexus A1×32-Poly2-5mm-50s-177-OZ32LP) with integrated fiber optics. Optical fibers had cores of 200 μm (0.39 NA, Pinnacle) or 50 μm (0.22 NA, Neuronexus). Mice were tethered to the recording system (Pinnacle: ADInstruments PowerLab and LabChart; Neuronexus: Tucker-Davis Technologies PZ5, RZ5D, and Synapse software), and LFPs were sampled at 1 Khz. All mice were habituated overnight to cylindrical recording chambers with access to food within the chamber and water *via* two chamber-mounted water bottles.

LFPs were continuously recorded throughout the 48-hour testing period. For analysis, we focused on lick-evoked neural activity and selected only those lick events that were not preceded by another lick for at least 6 seconds. This ensured that the observed BLA network dynamics reflected the onset of a new lick bout and were not confounded by lingering effects from earlier licks. For each qualifying lick bout, LFP segments spanning 6 seconds before and after the first lick were extracted for analysis.

To examine changes across the full spectral profile, we computed Morlet wavelet spectrograms for each trial using Python’s SciPy module. Spectrograms were baseline-referenced to their respective 3-second pre-lick baselines on a per-frequency basis. For band-limited analyses of identified frequency bands of interest, raw Morlet spectrograms were normalized per frequency by computing fold change from their respective 3-second baselines and then averaged across frequencies within each band, enabling assessment of relative power changes following lick initiation. Outlier removal was performed in two stages. At the sample level, transient artifacts (*e.g.,* mains noise and electrical spikes) were attenuated using Hampel filtering with a moving 600-sample window, replacing values exceeding 2.5-3.5 median absolute deviations from the local median with the corresponding median value. At the trial level, spectrograms were screened using power-based criteria: first, excessive amplitude deviations were detected using interquartile range and Tukey fence thresholds applied to baseline and band-limited power distributions; second, trials with total power or short-window (0-500 ms) power > 3 standard deviations from the group mean were excluded. Outlier indices were conservatively expanded (binary dilation) to remove neighboring contaminated samples, and remaining gaps were interpolated and filled using median replacement. In addition, for all frequency bands, trials were excluded if the final 4000 ms of the baseline-normalized recording had a mean > 10 a.u. or a peak-to-peak amplitude > 100 a.u., removing noisy trials. Because beta-band recordings were more susceptible to noise, an additional filtering step was applied: the 15-30 Hz band-limited time series was first extracted from the baseline-referenced spectrogram, subjected to additional Hampel filtering to suppress transient outliers, and then re-normalized to its own 3s pre-lick baseline, ensuring that residual artifacts did not inflate trial-averaged beta power.

#### Ex vivo

Mice were deeply anesthetized with isoflurane and decapitated using a rodent guillotine. Brains were rapidly extracted and immersed in ice-cold, oxygenated sucrose cutting solution containing (in mM): 150 sucrose, 33 NaCl, 25 NaHCO3, 2.5 KCl, 1.25 NaH2PO4, 1 CaCl2, 7 MgCl2, and 15 glucose (bubbled with 95% O2 / 5% CO2. Coronal sections (350 μm) were cut using a Leica VT1000S vibratome dedicated for live-tissue preparations. To eliminate hippocampal input, the BLA was resected from each slice in a petri dish and transferred to a 34 °C interface holding chamber containing oxygenated artificial cerebrospinal fluid (ACSF) containing (in mM): 126 NaCl, 10 glucose, 2 MgCl2, 2 CaCl2, 2.5 KCl, 1.25 NaH2PO4, 26 NaHCO3, 1.5 Na-pyruvate, and 1 L glutamine (300–310 mOsm). Slices were incubated for at least 1 hour before transfer to the interface recording chamber.

BLA extracellular field potential recordings were performed in the BLA using pulled borosilicate glass pipettes. Signals were amplified and digitized using LabChart (ADInstruments) at a sampling rate of 10 Khz and low-pass filtered at 3 Khz during acquisition. Gamma oscillations (30 - 60 Hz) were induced by continuous perfusion of modified ACSF containing elevated potassium (7.5 mM) and kainic acid (800 nM) at a rate of 1.799 mL/min. Each recording consisted of a 15-minute baseline period followed by a 15-minute drug treatment period.

For analysis, LFP data were extracted from the final 5 minutes of baseline and from minutes 3-8 of the drug treatment period to ensure full drug perfusion. For isoproterenol, which elicited more transient effects than norepinephrine, only minutes 3-5 of the treatment period were analyzed. This window was chosen to capture the peak of the drug-induced response, as our goal was to assess peak sustained gamma power modulation rather than effect duration. Time-frequency spectrograms were generated using the multitaper method (MNE-Python, time-bandwidth product = 4, 1s windows) to visualize gamma-band (30-60 Hz) activity. Spectrograms were aligned to the time of drug application (t = 0) and normalized to the baseline mean for qualitative comparison. Spectral decomposition for quantitative analysis was performed in Python using Welch’s method (SciPy) for analysis. Gamma power was similarly normalized to the mean baseline power for each slice.

### Drugs

All drugs used for bath application during slice electrophysiology were obtained from commercial suppliers. Kainate (Millipore Sigma Cat. # K0250), Noradrenaline bitartrate (Tocris Cat. # 5169), Isoproterenol (Tocris Cat. # 1747), WB4101 (Fisher Scientific Cat. # 11-101-3550), were prepared as concentrated stock solutions in DI water. WIN 55,212-2 (Tocris Cat. # 1038), m-3M3FBS (Tocris Cat. # 1941), FR 180204 (Tocris, Cat. # 3706), U0126 (Tocris, Cat. # 1144), and GF 109203X (Tocris, Cat. # 0741) were prepared as concentrated stock solutions in DMSO. All stock preparations were stored at −20 °C until use. On the day of recording, aliquots were diluted into ACSF to their final working concentrations immediately prior to bath application.

### Behavior

Mice were habituated overnight to cylindrical recording chambers equipped with *ad libitum* access to food and two chamber-mounted water bottles, each adapted from 15 mL conical tubes outfitted with pressure-sensitive lick spouts (modified from Frie and Khokhar, 2019^66^, Supplemental Figure 4). Lick events were recorded at 1 KHz using Arduino-controlled infrared beam-break lickometers. To detect the onset of new lick bouts for closed-loop optogenetic triggering, a bout was defined as any lick not preceded by another lick within the previous 6 seconds.

**Supplemental Figure 4.**
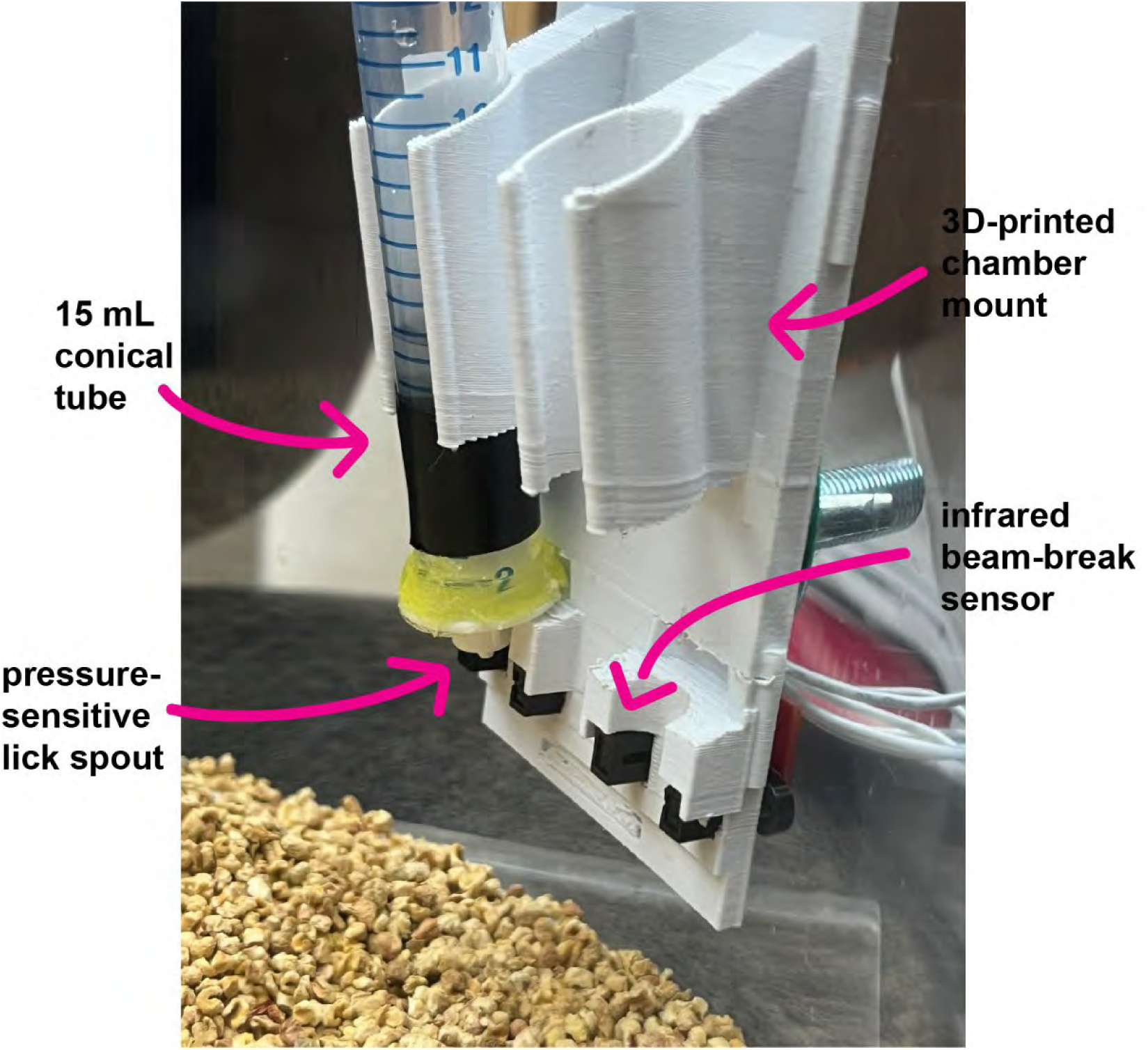
Chamber-mounted lickometer system. Representative image of the custom 3D-printed chamber-mounted bottle holder adapted from Frie and Khokhar (2019), designed to accommodate two modified 15 mL conical tubes equipped with pressure-sensitive lick spouts. Each spout is paired with an infrared beam-break sensor, with signals routed to an Arduino microcontroller for millisecond-resolution lick detection.

To distinguish between the two bottles, a thin strip of black electrical tape was affixed to one of the conical tubes (Supplemental Figure 4). During training, the left-right positions of the bottles were reversed every 24 hours to prevent side bias. Because we observed that mice often persisted with a single familiar spout following these reversals, lick spouts were wiped with DI water and a Kimwipe after each exchange, which encouraged mice to sample from both bottles. Behavioral testing/recording did not begin until each mouse had sampled from both bottles, with the first day on which both bottles were sampled designated as Day 1. During recording periods, bottles were left unperturbed except for daily contactless refilling through a hole at the top. In closed-loop optogenetic experiments, where bottles were filled with the same water from the same faucet at the same time, licks of the light-paired bottle triggered laser illumination of LC terminals in the BLA *via* a 640 nm DPSS benchtop laser (Laserglow Technologies), delivering ∼ 10 mW at 10 Hz with 60% pulse width. Importantly, stimulation persisted for the full duration of the licking bout plus an additional 3 seconds. Thus, if licking continued for *n* seconds, the total stimulation duration was *n* + 3 seconds, beginning with the first lick of the bout. No laser illumination was used for assessment of sucrose preference in PV knockdown experiments.

Behavior was assessed across two days. On day one, spontaneous bottle preference was quantified by counting lick bouts at each bottle over a 24-hour period. On day two, the previously preferred bottle was paired with laser illumination. Preference was calculated as the proportion of total lick bouts directed to each bottle, and aversion to the light-paired bottle was evaluated by comparing changes in preference across days.

Sucrose preference was also conducted in the same behavioral apparatus. Mice were habituated and individually housed in the cylindrical recording chambers overnight with *ad libitum* food and water access (except for food-deprived experiments where mice were dispensed a fixed amount of food pellets per day to maintain 80-90% bodyweight). On day one, both bottles were similarly filled with water, and lick events were recorded over 24 hours to establish baseline bottle preference. On day two, a 20% sucrose solution replaced the water in the previously non-preferred bottle. Bottle preference was quantified as the percentage of lick bouts directed to each bottle on both days, and sucrose preference was assessed accordingly.

### Statistical Methods

All frequentist statistical tests were conducted in Graphpad Prism 8 with test type and significance thresholds reported in the corresponding statistics tables. To complement these analyses and quantify relationships between neural activity and behavioral measures, we fit hierarchical Bayesian regression models in R (v4.4.3) using the brms package (v2.22.0), which implements Hamiltonian Monte Carlo sampling through Stan. To test the relationship between bout progression and bout duration during LC-BLA^NE^ activation, we fit a bayesian hierarchical model predicting log-transformed bout duration (with raw zero values replaced by 0.01s) from bout progression. We also tested the relationship between bout progression and IBI during LC-BLA^NE^ activation using a Bayesian hierarchical model predicting log-transformed IBI from bout progression. Both models included random intercepts and slopes for each mouse to account for between-subject variability in baseline bout duration and in the effect of bout progression. We placed weakly informative priors to regularize estimation: an exponential(5) prior on group-level standard deviations, a Student-t(−3, 0, 1) prior on fixed effects, and a normal (0, 1) prior on the residual standard deviation. For LC-BLA^NE^ experiments investigating relationships between behavior and BLA LFP activity, we modeled normalized fast gamma power (70-120 Hz) relative to baseline (centered by subtracting 1, such that baseline = 0) as a function of bout progression (bout index), including mouse as a grouping factor. In an additional model, log-transformed bout duration (with raw zero values replaced by 0.01s) was predicted from centered gamma power. Both models were initially specified with random intercepts and random slopes for each mouse to allow subject-specific deviations. However, the estimated between-subject slope variance was consistently near zero, and inclusion of this parameterization led to convergence difficulties. To improve stability and reflect the lack of meaningful slope heterogeneity, we simplified the random-effects structure to random intercepts only. Priors were chosen to be weakly informative but biologically plausible. For bout progression models, group-level intercept standard deviations were assigned Exponential(5) distributions, fixed-effect slopes were given Normal(0, 0.03) priors to constrain estimates near zero, and residual standard deviations were given Normal(0,1) priors. For bout duration models, group-level intercept standard deviations again followed Exponential(5), slopes were given Student-t(3,0,1) priors, and residual standard deviations Normal(0,1). In the PV DTA experiments, where identical models were attempted, we observed the same pattern of negligible between-subject slope variability. Accordingly, final models were fit with random intercepts only. All models were estimated using four to six independent chains with 4000 total iterations per chain, of which half were reserved for warmup. To ensure robust sampling, the step size target acceptance was set between 0.95 and 0.999, and the maximum tree depth was left at the default of 10 for all models except for the two predicting log-transformed bout durations or IBIs as a function of bout progression in LC-BLA^NE^ experiments, where it was set to 12. All Rhat values were <1.01 and effective sample sizes (ESS) were > 1000. Inference was based on posterior means and 95% credible intervals, and posterior probabilities for directional effects were computed directly from posterior draws. Model fit was further quantified using Bayesian *R*^2^.

### Histology & Microscopy

#### Perfusion and Sectioning

Prior to brain extraction, mice were deeply anesthetized with ketamine/xylazine cocktail and transcardially perfused with PBS until the exposed liver displayed significant lightening, followed by 12.5 mL 4% paraformaldehyde in PBS. Extracted brains were left in 4% PFA in PBS overnight at 4 °C and kept in PBS only until sectioning. Coronal sections (50 μm) were cut using a Leica VT1000S vibratome and stored in 24-well plates at 4 °C.

#### Immunofluorescence

Free-floating sections were washed 3 times in a PBS solution containing 0.2% Triton X-100 (PBS-Tx) for 5 minutes each. Following washes, sections were incubated in blocking solution (10% normal goat serum in PBS-Tx) for 1 hour at room temperature. Sections were then incubated in primary antibody solution containing antibodies against PV (1:2000 in blocking solution, Guinea Pig anti-PV, Synaptic Systems cat. # 195 004) overnight at 4 °C. The following day, sections were washed 3 times for 5 minutes each in PBS-Tx and placed in Goat anti-Guinea Pig (AF647) secondary antibody solution (Thermo Fisher cat. # A-21450) for 2h at room temperature in the dark. Note that all incubation steps were performed with continuous gentle agitation on a benchtop rocker. Following secondary incubation, sections were washed 3 times in PBS in the dark, mounted on microscope slides, and allowed to dry in the dark until matte. Finally, mounted slides were coverslipped using Prolong Gold Antifade Mountant (Thermo Fisher cat. # P36930).

#### Dual Immunofluorescence and In Situ Hybridization

To visualize mRNA and protein expression in the same tissue sections for representative *ADRA1A* expression in Dbh-Cre and Dbh-Cre^α1AKO^ mice, we adapted the ACD Bio RNAScope Multiplex Fluorescent v2 Assay for use with immunofluorescence in free-floating, PFA-fixed mouse brain sections. Mice were perfused with 4% PFA in PBS, post-fixed overnight, and cut into 50 μm coronal sections as described above. RNAScope target retrieval was performed by incubating tissue in pre-heated Target Retrieval Agent (ACD Bio) within a tabletop steamer (Oster) for 5 minutes. Probes (Cat. # 408611-C2, 413561-C3) were prepared using a 1:50 ratio of labeled probes (C1-C4) to the negative control or C1 backbone probe. Tissue was left to incubate in the final probe mixture for 2 hours at 40 °C in a benchtop incubator. Signal amplification steps (AMP1-3, HRP-conjugated channel-specific reagents) and TSA fluorophore addition were performed sequentially at 40 °C following the ACD Bio RNAScope Multiplex Fluorescent v2 protocol with 2-minute washes in 1X Wash Buffer between steps. HRP blocking was included between fluorophore channels, and tissue was protected from light after the first fluorophore incubation.

Following RNAScope staining, tissue was washed 3 times in 0.2% PBS-Tx (5 minutes each) and incubated in blocking solution (10% normal goat serum in 0.2% PBS-Tx) for 1 hour at room temperature. Sections were next incubated in Guinea Pig anti-PV primary antibody solution (1:2000 in blocking solution, Synaptic Systems cat. # 195 004) for 2 hours at room temperature. Following primary incubation, sections were washed 3 times with 0.2% PBS-Tx (5 minutes each) and placed in Goat anti-Guinea Pig (AF647) secondary antibody solution (Thermo Fisher cat. # A-21450) for 2h at room temperature in the dark. Sections were finally washed 3 times with PBS (5 minutes each), incubated with DAPI (ACD Bio cat. # 320858) solution for 30 seconds and mounted onto non-charged glass slides with ProLong Gold Antifade Mountant (Thermo Fisher cat. # P36930). Note that all incubation steps were performed with continuous gentle agitation on a benchtop rocker. Further, all tissue used in this protocol was sectioned within 1-2 days prior to staining to prevent damage during target retrieval. Importantly, this protocol avoids protease treatment to preserve protein antigenicity and was optimized for thick (50 μm) vibratome sections to maintain free-floating tissue integrity throughout the RNAScope workflow.

#### Microscopy and Image Deconvolution

Widefield fluorescence images were acquired using a Nikon TE2000E inverted microscope fitted with Thermo Photometrics X1 Camera and EXFO X-CITE 120XL metal halide illuminator. Basic image post-processing (*e.g.,* color) was performed in Fiji. High magnification (60x) fluorescence images were deconvolved using Deconwolf^67^, an open-source image deconvolution software optimized for widefield microscopy data. Deconvolution was performed using a theoretical point spread function generated in Fiji using the PSF Generator plugin, parameterized based on imaging system metadata (*e.g.,* numerical aperture, emission wavelength, refractive index). Following deconvolution, immunofluorescence and *in situ* hybridization image z-stacks were flattened using the “Extended Depth of Focus Sobel Projection” plugin from the CLIJ2 suite and maximum intensity projection function in Fiji, respectively.

## Statistics Tables

**Statistics Table 1.**
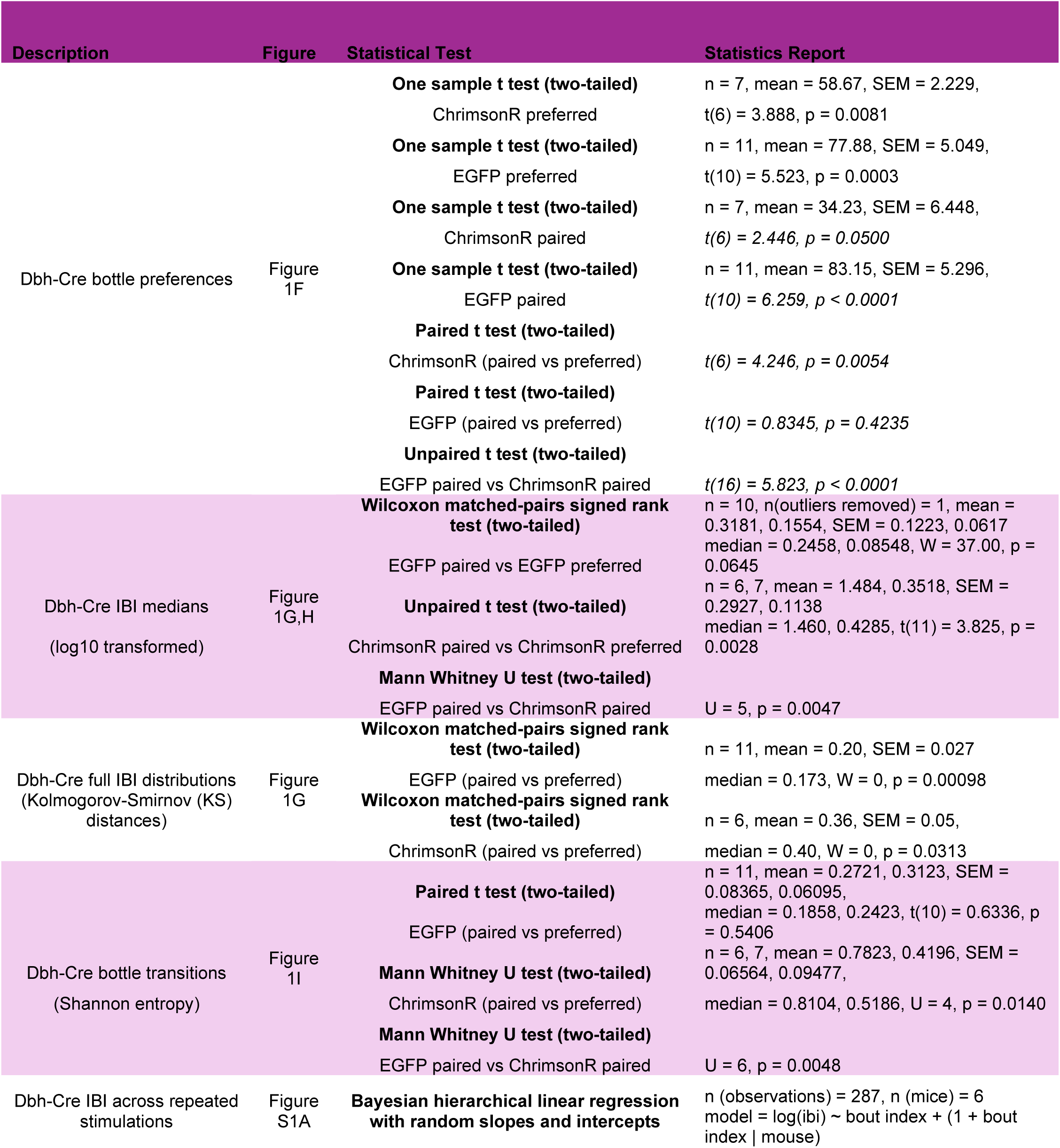
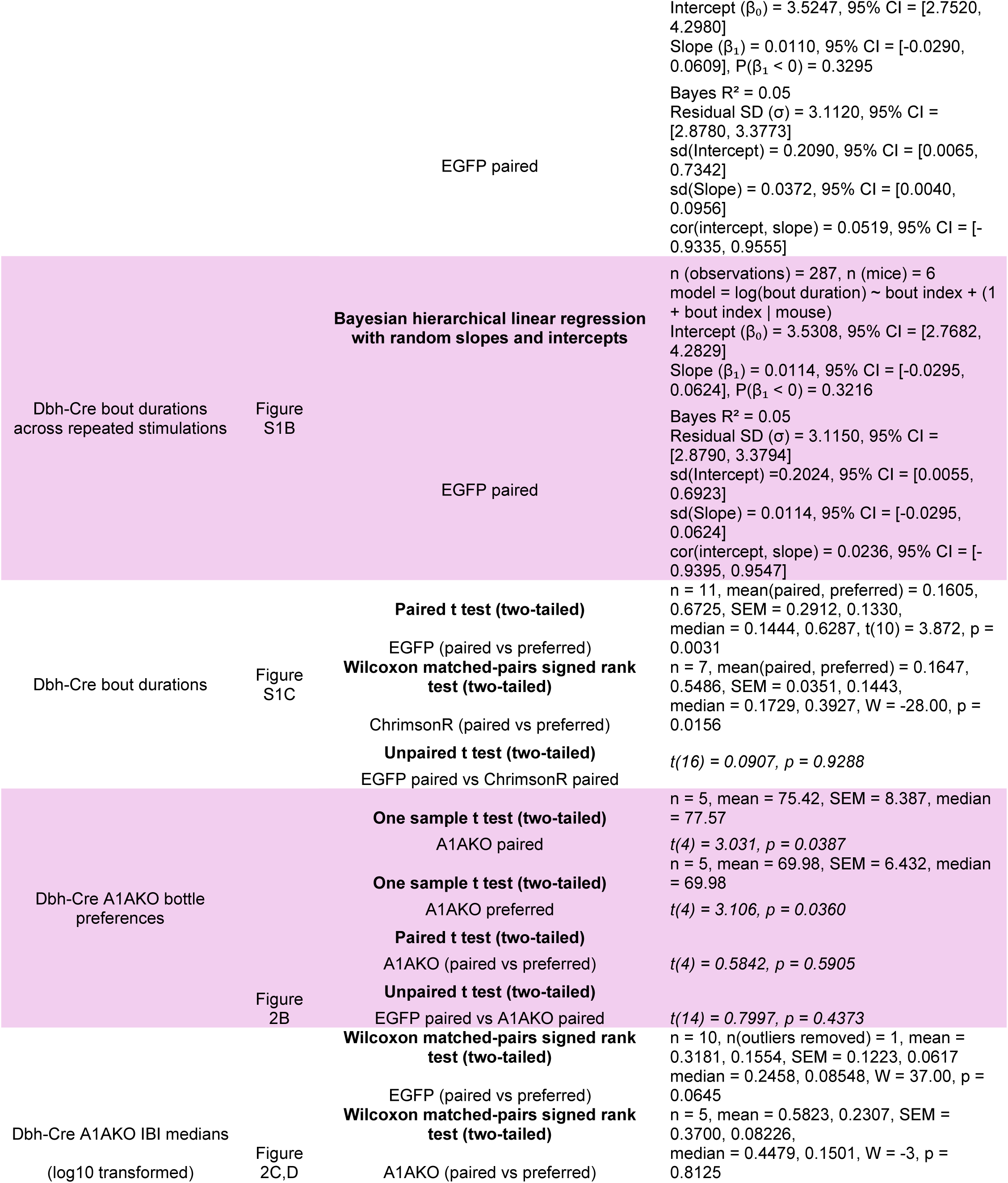
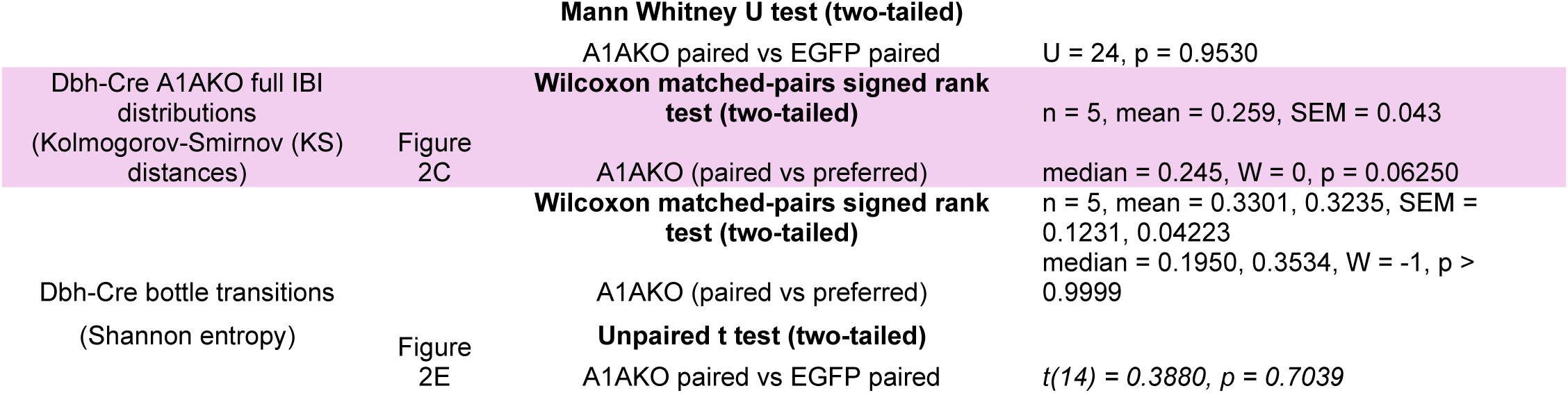

**Statistics Table 2.**
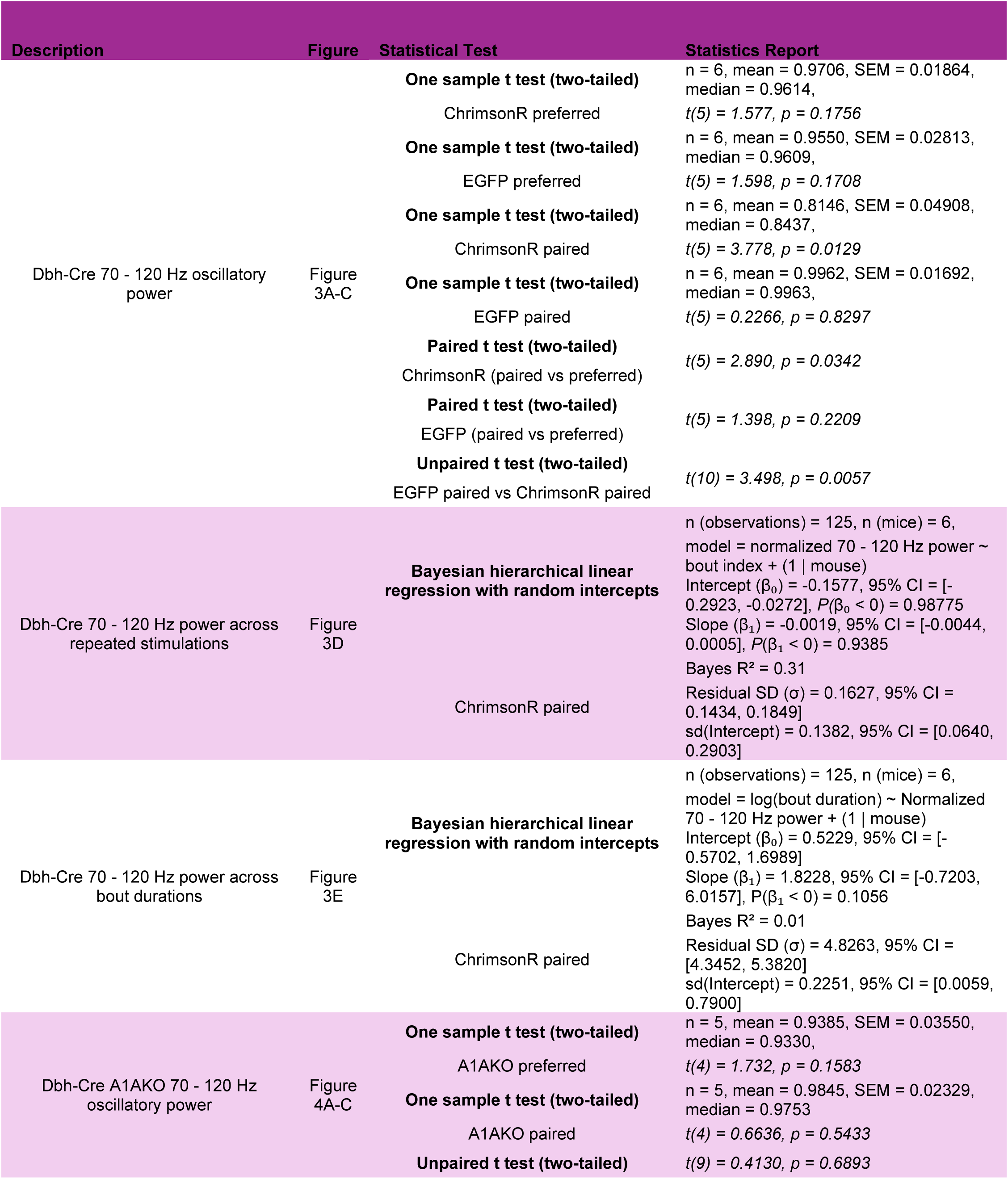
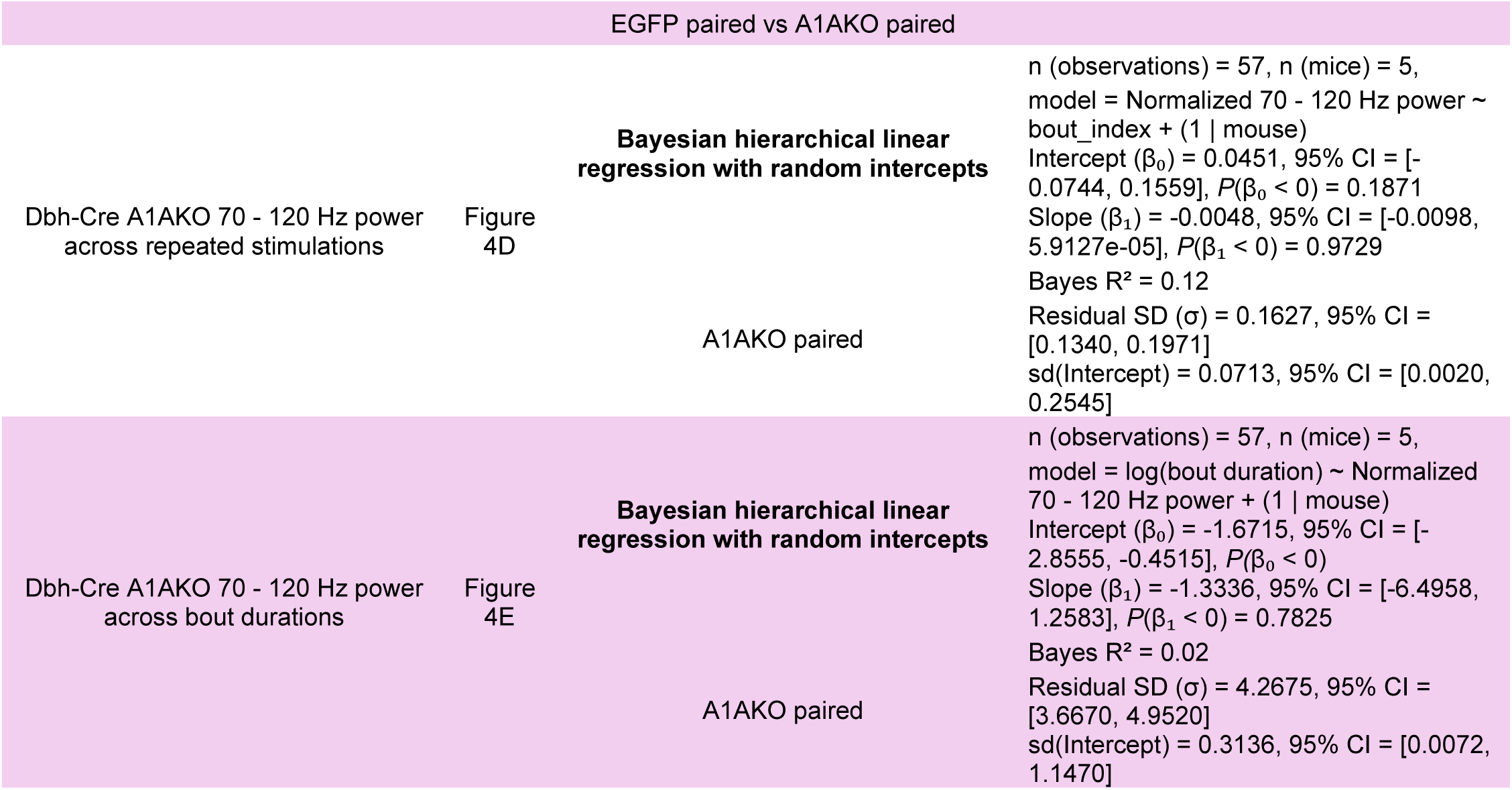

**Statistics Table 3.**
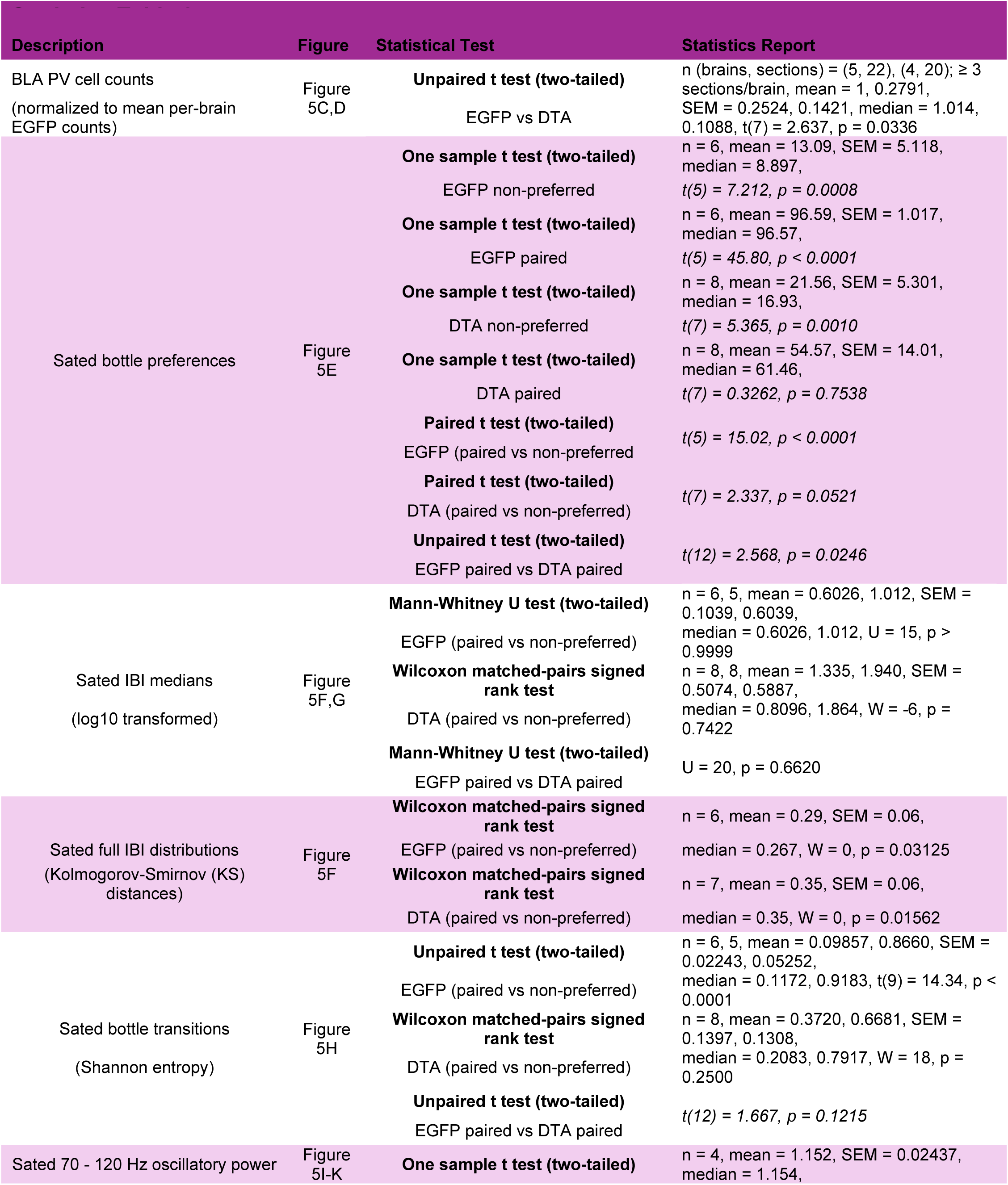
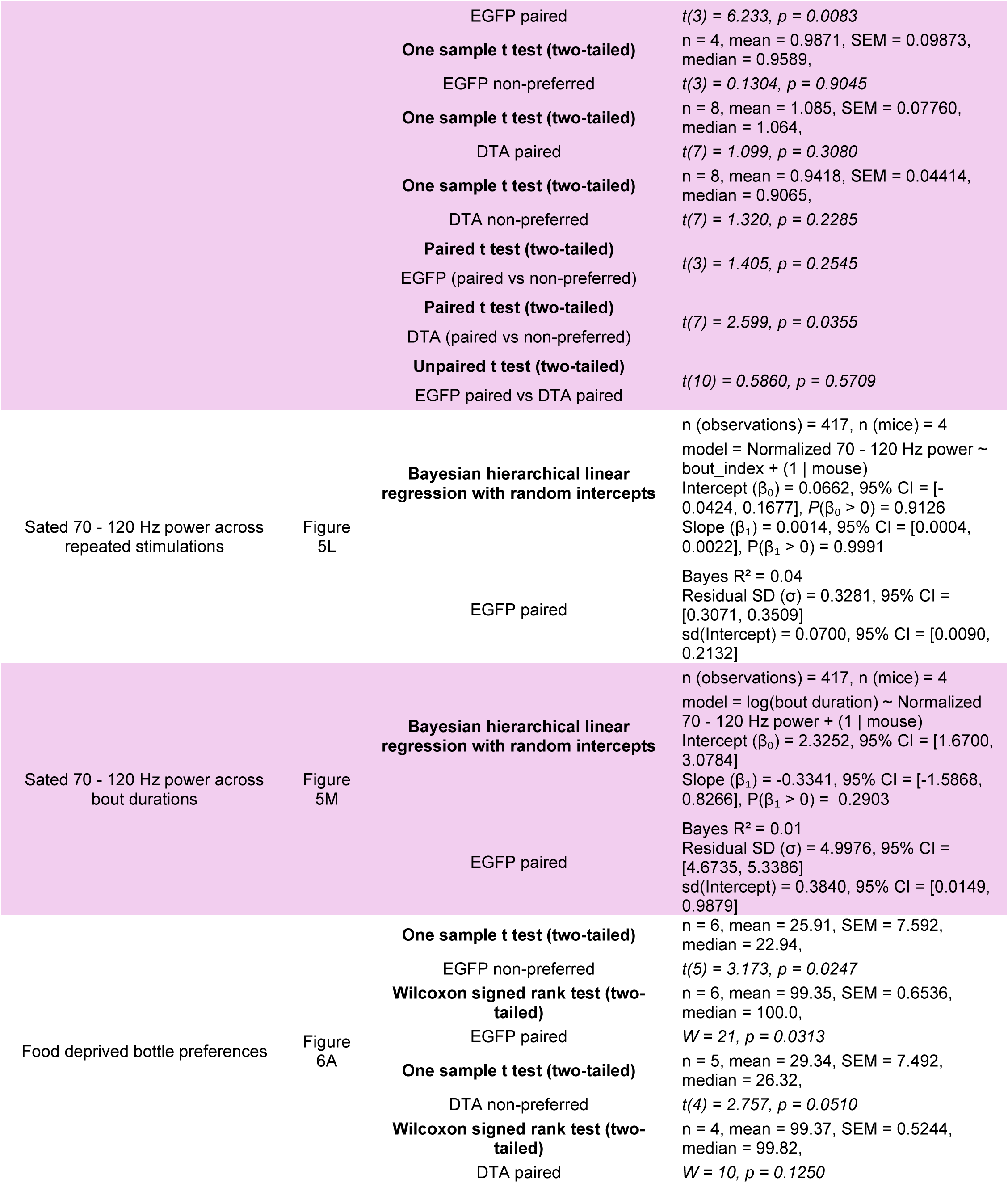
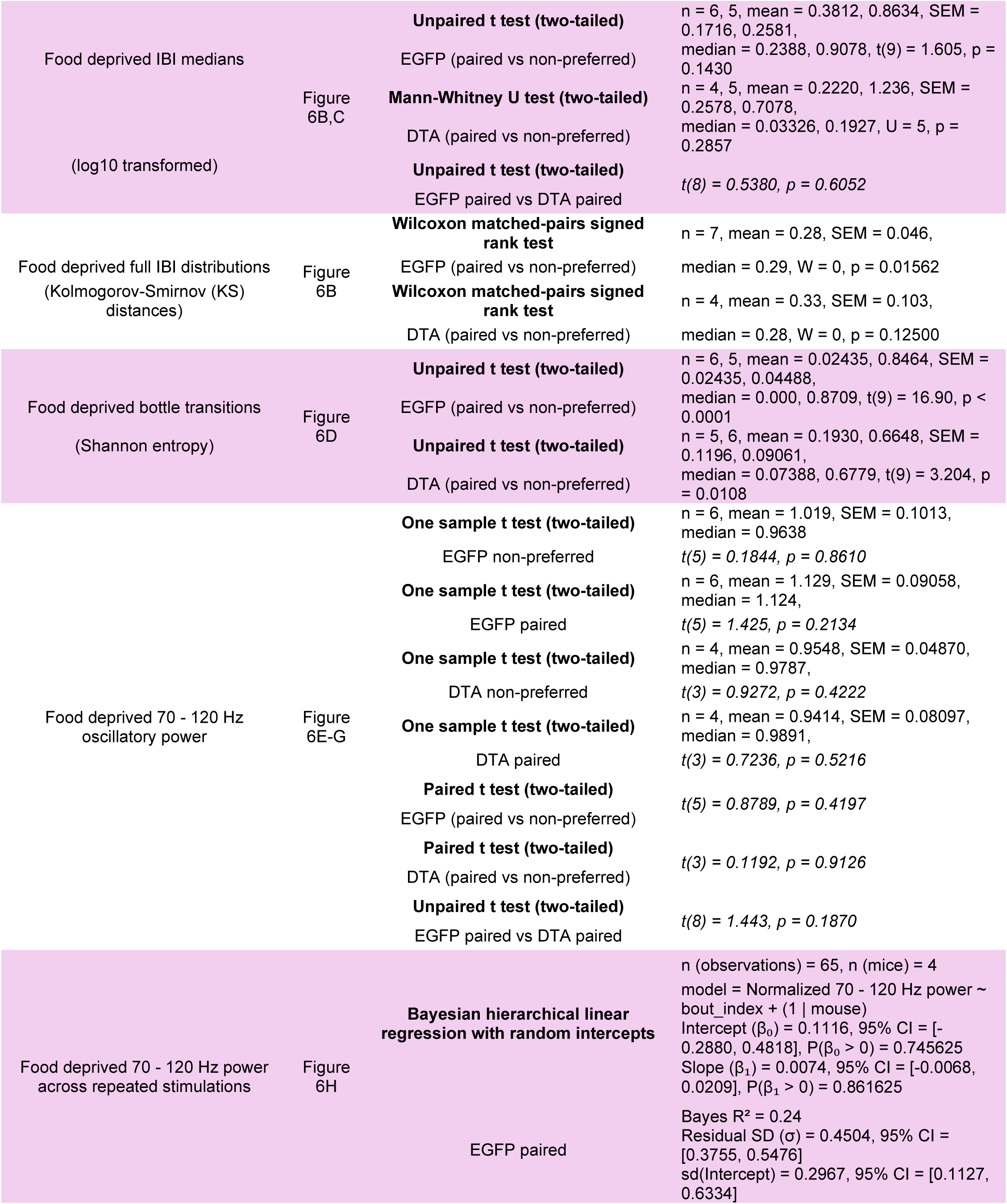
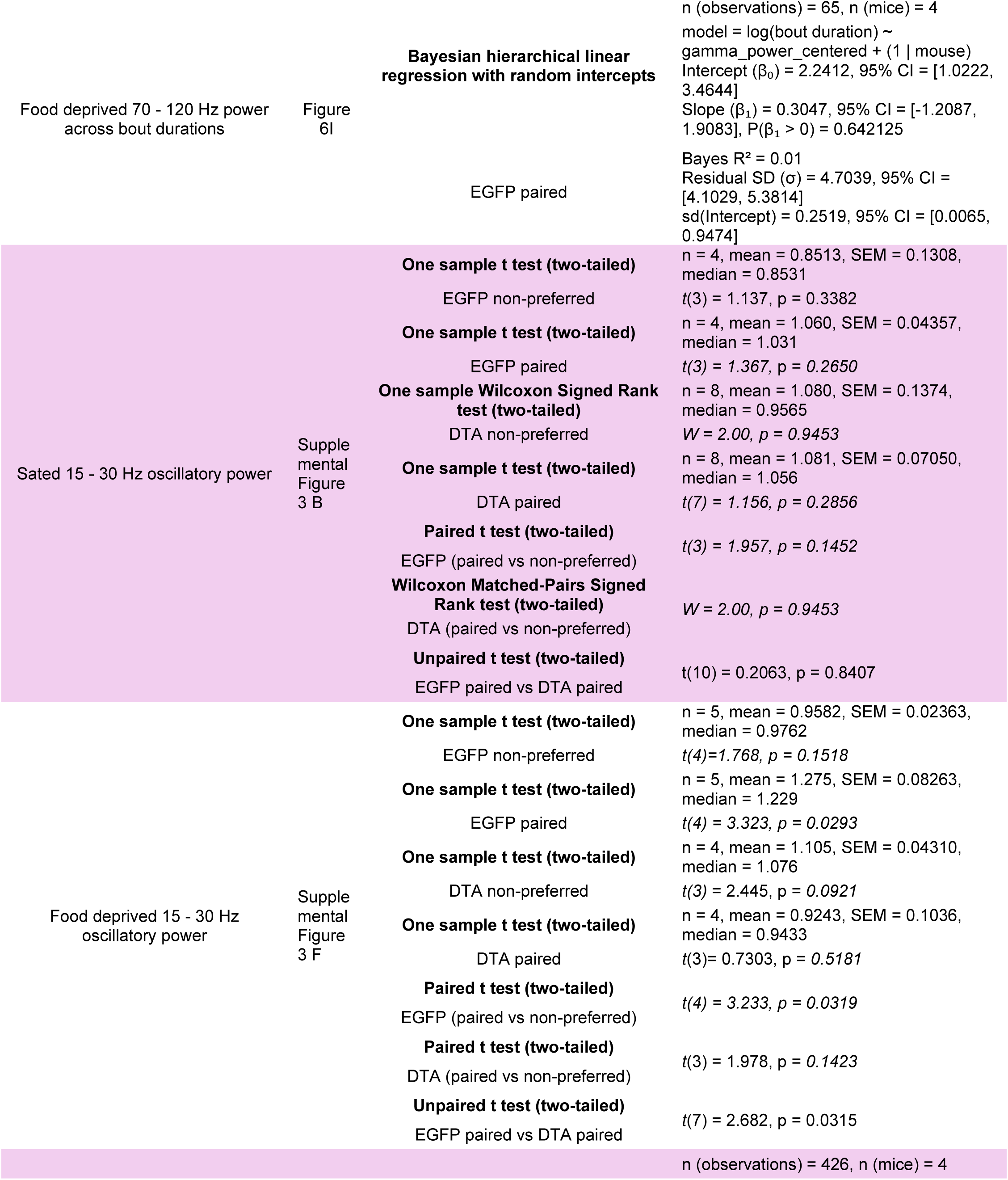
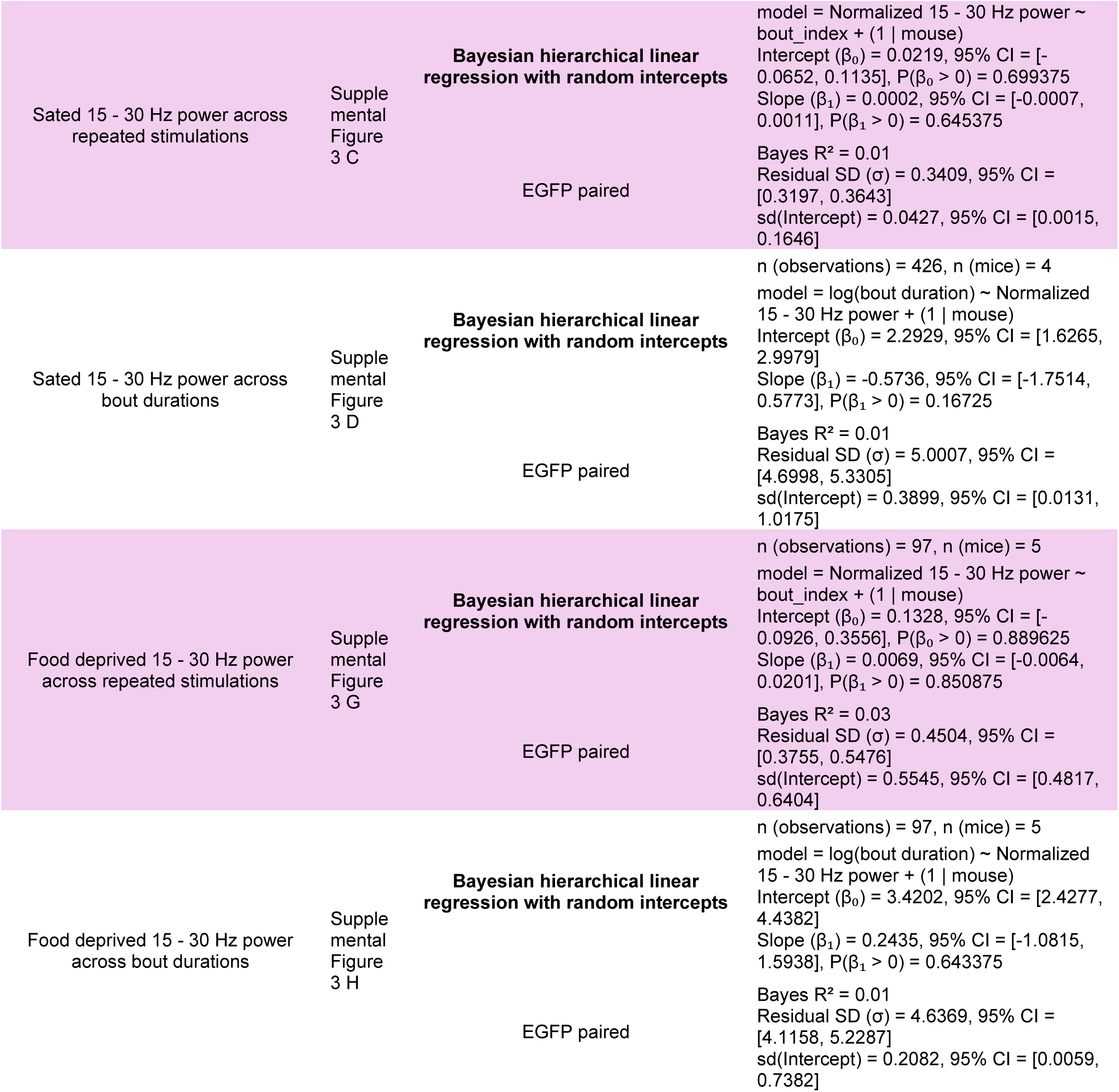

## References

1. Tye, K. M. Neural circuit motifs in valence processing. Neuron 100, 436–452 (2018).

2. Ben-Zion, Z. et al. Neural responsivity to reward versus punishment shortly after trauma predicts long-term development of posttraumatic stress symptoms. Biol. Psychiatry Cogn. Neurosci. Neuroimaging 7, 150–161 (2022).

3. Medeiros, G. C. et al. Positive and negative valence systems in major depression have distinct clinical features, response to antidepressants, and relationships with immunomarkers. Depress. Anxiety 37, 771–783 (2020).

4. Hu, S. et al. Correlation between suicidal ideation and emotional memory in adolescents with depressive disorder. Sci. Rep. 12, 5470 (2022).

5. Smith, P. N., Cukrowicz, K. C., Poindexter, E. K., Hobson, V. & Cohen, L. M. The acquired capability for suicide: a comparison of suicide attempters, suicide ideators, and non-suicidal controls. Depress. Anxiety 27, 871–877 (2010).

6. Wen, A., LeMoult, J., McCabe, R. & Yoon, K. L. Affective flexibility and generalized anxiety disorder: valence-specific shifting difficulties. Anxiety Stress Coping 32, 581–593 (2019).

7. Antonoudiou, P. et al. Biased information routing through the basolateral amygdala, altered valence processing, and impaired affective states associated with psychiatric illnesses. Biol. Psychiatry 97, 764–774 (2025).

8. Brown, S. & Schafer, E. A. XI. An investigation into the functions of the occipital and temporal lobes of the monkey’s brain. Philos. Trans. R. Soc. Lond. 179, 303–327 (1888).

9. Weiskrantz, L. Behavioral changes associated with ablation of the amygdaloid complex in monkeys. J. Comp. Physiol. Psychol. 49, 381–391 (1956).

10. ‘Psychic blindness’ and other symptoms following bilateral temporal lobectomy in Rhesus monkeys. American Journal of Physiology, 119, (1937).

11. Paton, J. J., Belova, M. A., Morrison, S. E. & Salzman, C. D. The primate amygdala represents the positive and negative value of visual stimuli during learning. Nature 439, 865–870 (2006).

12. Zhang, X. & Li, B. Population coding of valence in the basolateral amygdala. Nat. Commun. 9, 5195 (2018).

13. Beyeler, A. et al. Divergent routing of positive and negative information from the amygdala during memory retrieval. Neuron 90, 348–361 (2016).

14. Moryś, J. et al. Relationship of calcium-binding protein containing neurons and projection neurons in the rat basolateral amygdala. Neurosci. Lett. 259, 91–94 (1999).

15. Redondo, R. L. et al. Bidirectional switch of the valence associated with a hippocampal contextual memory engram. Nature 513, 426–430 (2014).

16. Kim, J., Pignatelli, M., Xu, S., Itohara, S. & Tonegawa, S. Antagonistic negative and positive neurons of the basolateral amygdala. Nat. Neurosci. 19, 1636–1646 (2016).

17. Zhang, X. et al. Genetically identified amygdala-striatal circuits for valence-specific behaviors. Nat. Neurosci. 24, 1586–1600 (2021).

18. Beyeler, A. et al. Organization of valence-encoding and projection-defined neurons in the basolateral amygdala. Cell Rep. 22, 905–918 (2018).

19. Dwyer, D. M. Lesions of the basolateral, but not central, amygdala impair flavour-taste learning based on fructose or quinine reinforcers. Behav. Brain Res. 220, 349–353 (2011).

20. Shen, C.-J. et al. Cannabinoid CB1 receptors in the amygdalar cholecystokinin glutamatergic afferents to nucleus accumbens modulate depressive-like behavior. Nat. Med. 25, 337–349 (2019).

21. Namburi, P. et al. A circuit mechanism for differentiating positive and negative associations. Nature 520, 675–678 (2015).

22. Namburi, P., Al-Hasani, R., Calhoon, G. G., Bruchas, M. R. & Tye, K. M. Architectural representation of valence in the limbic system. Neuropsychopharmacology 41, 1697–1715 (2016).

23. Li, H. et al. Neurotensin orchestrates valence assignment in the amygdala. Nature 608, 586–592 (2022).

24. Omoluabi, T., Power, K. D., Sepahvand, T. & Yuan, Q. Phasic and tonic locus coeruleus stimulation associated valence learning engages distinct adrenoceptors in the rat basolateral amygdala. Front. Cell. Neurosci. 16, 886803 (2022).

25. Ghosh, A. et al. Locus coeruleus activation patterns differentially modulate odor discrimination learning and odor valence in rats. Cereb. Cortex Commun. 2, tgab026 (2021).

26. McCall, J. G. et al. CRH engagement of the locus coeruleus noradrenergic system mediates stress-induced anxiety. Neuron 87, 605–620 (2015).

27. McCall, J. G. et al. Locus coeruleus to basolateral amygdala noradrenergic projections promote anxiety-like behavior. Elife 6, (2017).

28. Bigot, M. et al. Disrupted basolateral amygdala circuits supports negative valence bias in depressive states. Transl. Psychiatry 14, 382 (2024).

29. Day, H. E. W., Campeau, S., Watson, S. J., Jr & Akil, H. Distribution of α1a-, α1b- and α1d-adrenergic receptor mRNA in the rat brain and spinal cord. J. Chem. Neuroanat. 13, 115–139 (1997).

30. Fu, X. et al. Gq neuromodulation of BLA parvalbumin interneurons induces burst firing and mediates fear-associated network and behavioral state transition in mice. Nat. Commun. 13, 1290 (2022).

31. Colmers, P. L. W. et al. Loss of PV interneurons in the BLA may contribute to altered network and behavioral states in chronically epileptic mice. eNeuro 12, ENEURO.0482–23.2024 (2025).

32. Antonoudiou, P. et al. Allopregnanolone mediates affective switching through modulation of oscillatory states in the basolateral amygdala. Biol. Psychiatry 91, 283–293 (2022).

33. Dwarakanath, A. et al. Bistability of prefrontal states gates access to consciousness. Neuron 111, 1666–1683.e4 (2023).

34. Pinotsis, D. A., Fridman, G. & Miller, E. K. Cytoelectric coupling: Electric fields sculpt neural activity and ‘tune’ the brain’s infrastructure. Prog. Neurobiol. 226, 102465 (2023).

35. Umbach, G., Tan, R., Jacobs, J., Pfeiffer, B. E. & Lega, B. Flexibility of functional neuronal assemblies supports human memory. Nat. Commun. 13, 6162 (2022).

36. Davis, P., Zaki, Y., Maguire, J. & Reijmers, L. G. Cellular and oscillatory substrates of fear extinction learning. Nat. Neurosci. 20, 1624–1633 (2017).

37. Ozawa, M. et al. Experience-dependent resonance in amygdalo-cortical circuits supports fear memory retrieval following extinction. Nat. Commun. 11, 4358 (2020).

38. Stujenske, J. M. et al. Prelimbic cortex drives discrimination of non-aversion via amygdala somatostatin interneurons. Neuron 110, 2258–2267.e11 (2022).

39. Stujenske, J. M., Likhtik, E., Topiwala, M. A. & Gordon, J. A. Fear and safety engage competing patterns of theta-gamma coupling in the basolateral amygdala. Neuron 83, 919–933 (2014).

40. Likhtik, E., Stujenske, J. M., Topiwala, M. A., Harris, A. Z. & Gordon, J. A. Prefrontal entrainment of amygdala activity signals safety in learned fear and innate anxiety. Nat. Neurosci. 17, 106–113 (2014).

41. Amaya, K. A., Teboul, E., Weiss, G. L., Antonoudiou, P. & Maguire, J. L. Basolateral amygdala parvalbumin interneurons coordinate oscillations to drive reward behaviors. Curr. Biol. 34, 1561–1568.e4 (2024).

42. Ni, J. et al. Gamma-rhythmic gain modulation. Neuron 92, 240–251 (2016).

43. Sohal, V. S., Zhang, F., Yizhar, O. & Deisseroth, K. Parvalbumin neurons and gamma rhythms enhance cortical circuit performance. Nature 459, 698–702 (2009).

44. Womelsdorf, T., Valiante, T. A., Sahin, N. T., Miller, K. J. & Tiesinga, P. Dynamic circuit motifs underlying rhythmic gain control, gating and integration. Nat. Neurosci. 17, 1031–1039 (2014).

45. Siegle, J. H., Pritchett, D. L. & Moore, C. I. Gamma-range synchronization of fast-spiking interneurons can enhance detection of tactile stimuli. Nat. Neurosci. 17, 1371–1379 (2014).

46. Kanta, V., Pare, D. & Headley, D. B. Closed-loop control of gamma oscillations in the amygdala demonstrates their role in spatial memory consolidation. Nat. Commun. 10, 3970 (2019).

47. Park, A. J. et al. Reset of hippocampal-prefrontal circuitry facilitates learning. Nature 591, 615–619 (2021).

48. Rokosh, D. G. & Simpson, P. C. Knockout of the alpha 1A/C-adrenergic receptor subtype: the alpha 1A/C is expressed in resistance arteries and is required to maintain arterial blood pressure. Proc. Natl. Acad. Sci. U. S. A. 99, 9474–9479 (2002).

49. Hájos, N. Interneuron types and their circuits in the basolateral amygdala. Front. Neural Circuits 15, 687257 (2021).

50. Perumal, M. B. & Sah, P. Inhibitory circuits in the basolateral amygdala in aversive learning and memory. Front. Neural Circuits 15, 633235 (2021).

51. Stone, B. T. et al. Early life stress impairs VTA coordination of BLA network and behavioral states. J. Neurosci. 45, e0088242025 (2025).

52. Pappenheimer, A. M., Jr. Diphtheria toxin. Annu. Rev. Biochem. 46, 69–94 (1977).

53. Hooper, A., Fuller, P. M. & Maguire, J. Hippocampal corticotropin-releasing hormone neurons support recognition memory and modulate hippocampal excitability. PLoS One 13, e0191363 (2018).

54. Tye, K. M. & Janak, P. H. Amygdala neurons differentially encode motivation and reinforcement. J. Neurosci. 27, 3937–3945 (2007).

55. Stuber, G. D. et al. Excitatory transmission from the amygdala to nucleus accumbens facilitates reward seeking. Nature 475, 377–380 (2011).

56. Copik, A. J. et al. Isoproterenol acts as a biased agonist of the alpha-1A-adrenoceptor that selectively activates the MAPK/ERK pathway. PLoS One 10, e0115701 (2015).

57. Mizuno, N. & Itoh, H. Functions and regulatory mechanisms of Gq-signaling pathways. Neurosignals 17, 42–54 (2009).

58. Freund, T. F. & Katona, I. Perisomatic inhibition. Neuron 56, 33–42 (2007).

59. Jain, R., Watson, U., Vasudevan, L. & Saini, D. K. ERK activation pathways downstream of GPCRs. Int. Rev. Cell Mol. Biol. 338, 79–109 (2018).

60. Bae, Y.-S. et al. Identification of a compound that directly stimulates phospholipase C activity. Mol. Pharmacol. 63, 1043–1050 (2003).

61. Harris, C., Weiss, G. L., Di, S. & Tasker, J. G. Cell signaling dependence of rapid glucocorticoid-induced endocannabinoid synthesis in hypothalamic neuroendocrine cells. Neurobiol. Stress 10, 100158 (2019).

62. Weeber, E. J. et al. A role for the beta isoform of protein kinase C in fear conditioning. The Journal of neuroscience : the official journal of the Society for Neuroscience 20, (2000).

63. Fujita, S., Yoshida, S., Matsuki, T., Jaiswal, M. K. & Seki, K. The α1-adrenergic receptors in the amygdala regulate the induction of learned despair through protein kinase C-beta signaling. Behav. Pharmacol. 32, 73–85 (2021).

64. Feng, F. et al. Gamma oscillations in the basolateral amygdala: Biophysical mechanisms and computational consequences. eNeuro 6, ENEURO.0388–18.2018 (2019).

65. Schoenbaum, G., Chiba, A. A. & Gallagher, M. Neural encoding in orbitofrontal cortex and basolateral amygdala during olfactory discrimination learning. J. Neurosci. 19, 1876–1884 (1999).

65. Frie, J. A. & Khokhar, J. Y. An open source automated two-bottle choice test apparatus for rats. HardwareX 5, e00061 (2019).

67. Wernersson, E. et al. Deconwolf enables high-performance deconvolution of widefield fluorescence microscopy images. Nat. Methods 21, 1245–1256 (2024).

